# ANDROGENS PROTECT ILC2S FROM FUNCTIONAL SUPPRESSION DURING INFLUENZA VIRUS INFECTION

**DOI:** 10.1101/2025.01.23.634583

**Authors:** Sapana Kadel, Reegan A. J. Miller, Anna Karlik, Sean Turner, Abigael P. Williams, Erola Ainsua-Enrich, Ibrahim Hatipoglu, Harini Bagavant, José Alberola-Ila, Joshua W. Garton, Richard Pelikan, Susan Kovats

## Abstract

Biological sex differences in morbidity upon influenza A virus (IAV) infection are linked to stronger IFN-centered immune responses in females, yet the regulatory role of sex hormone receptors in immune cell subsets is incompletely understood. Lung-resident group 2 innate lymphoid cells (ILC2s) express notably high levels of androgen receptors (AR). In IAV infection, ILC2s produce type 2 cytokines and facilitate tissue repair, but they also may be functionally suppressed by type 1 cytokines. Here we report sex differences in the magnitude of lung ILC2 functional suppression at the peak of sublethal IAV infection. Relative to males, ILC2s in females show attenuated proliferation, decreased propensity for IL-5 and amphiregulin production and reduced expression of GATA3 and IL-33R, features supported by divergent transcriptomes. Equivalent inflammatory cytokine levels and viral load suggested sex differences in ILC2-intrinsic factors. Indeed, naïve female ILC2s showed elevated IFNGR expression and higher phospho-STAT1 levels following IFNγ stimulation, and lymphocyte-restricted STAT1 deficiency reversed IAV-induced suppression of female ILC2s. Lymphocyte-restricted AR deficiency or loss of androgens via orchiectomy led to increased IFNGR expression and suppression of male ILC2s. These data support the hypothesis that intrinsic AR activity regulates IFNGR-STAT1 signaling pathways to preserve canonical ILC2 function in males during IAV infection.

## Introduction

Women in their reproductive years often experience greater morbidity and mortality upon influenza virus infection, while severe disease in the elderly is more frequent in men, particularly in pandemic influenza outbreaks (1,2). This biological sex disparity has been linked to stronger IFN-centered innate and adaptive antiviral immune responses in women, which can lead to increased immunopathogenesis (2,3). In murine (C57BL/6) models of sublethal influenza A (IAV) infection, young females also showed increased morbidity secondary to greater type I inflammation, while decreased testosterone levels in older males promoted severe disease (4,5). Sex hormones have been implicated in the sex disparity observed in influenza infection (1,2), and immune cell subsets express sex hormone receptors to varying degrees, consistent with their regulation by androgens and estrogens (3). In infection and autoimmune models, estrogens are linked to enhanced IFN responses in females, while higher levels of androgens in males attenuate IFN-centered inflammation (2,3). Although sex-disparate responses of multiple immune cell types account for observed sex differences in IAV outcomes, few studies have linked sex hormone receptor on immune cells to their functional responses in females and males. Notably, murine group 2 innate lymphoid cells (ILC2s) express high levels of androgen receptor (*Ar*) mRNA compared to other lymphocytes (6–9). Herein, we use a murine model of IAV infection to dissect sex differences in ILC2 function.

ILCs are innate tissue-resident counterparts of T cells, and multiple ILC subsets play diverse roles to shape the inflammatory environment during viral infection. ILCs are subdivided into group 1 [ILC1 and natural killer cells (NKs)], group 2 (ILC2) and group 3 (ILC3) subsets based on their transcription factor and effector cytokine profiles. In the lung, ILC2s (GATA3^hi^) are poised to produce type 2 cytokines in response to alarmins such as IL-33 released from damaged epithelium (10,11). Early studies support the hypothesis that ILC2s are beneficial during respiratory virus infection as they promote viral clearance and facilitate tissue repair through their production of IL-5 and the EGF receptor-binding amphiregulin (AREG) (12,13). Later studies showed that the type I environment, involving interferons (IFNs) and other pro-inflammatory cytokines, suppresses ILC2 functional responses and promotes ILC2 to ILC1 plasticity during respiratory virus infection (14–22). Prior reports showed that IFNγ acting via the IFNγ receptor (IFNGR) results in reduced ILC2 proliferation and IL-5 production in a pathway dependent on STAT1 (14,16). STAT1 signaling also promotes ILC1 and restrains ILC2 responses upon respiratory syncytial virus infection (20). However, sex differences in IFNGR or STAT1 signaling in ILC2s during IAV infection have not been reported.

We and others have identified sex differences in the numbers and functional responses of lung resident ILC2s (6,7,23–25). Male sex and androgen receptor (AR) activity dampen ILC2 numbers and function in homeostasis and allergy models (6,7,23), but we lacked information about sex differences in ILC2 function during type I inflammation elicited by respiratory viruses. Elegant studies have identified the mechanistic roles of sex hormone receptor signaling in diverse immune cells and lung epithelium in homeostasis and respiratory virus infection [reviewed in (26,27)]. AR is a nuclear hormone receptor capable of mediating epigenetic chromatin changes that alter accessibility of regulated genes (28). Yet we do not fully understand the molecular and cellular mechanisms by which AR activity regulates ILC2 functional responses during type I inflammation.

Herein, we tested the hypothesis that sex differences in ILC2 numbers, plasticity and functional responses occur during murine respiratory virus infection. We show that the type I inflammation elicited by a sublethal IAV infection preferentially suppressed lung-resident ILC2s in females. Relative to males, ILC2s in females show attenuated proliferation, decreased propensity for IL-5 and AREG production and reduced expression of GATA3 and IL-33R. These functional sex differences were supported by single cell RNA sequencing data showing distinct transcriptome clustering in female and male ILC2s. The inflammatory cytokine levels and viral load were equivalent in both sexes, suggesting sex differences in ILC2-intrinsic factors. Indeed, naïve female ILC2s showed higher expression of IFNGR and higher phospho-STAT1 levels following stimulation by IFNγ, and lymphocyte-restricted STAT1 deficiency reversed suppression of female ILC2s during infection. Lung resident ILC2s express high levels of AR protein relative to other lymphocytes, and use of IL-7R-Cre and CD4-Cre drivers of *Ar* deletion showed that AR deficiency in IL-7R^+^ lymphocytes, yet not in T cells, led to increased IFNGR expression and elevated suppression in male ILC2s. Depletion of androgens by orchiectomy abolished the sex differences in multiple parameters including IFNGR expression, leading to ILC2 suppression in orchiectomized males that was comparable to females post-infection. Our data support the hypothesis that the AR and IFNGR-STAT1 signaling pathways mediate ILC2-intrinsic sex differences during influenza virus infection.

## Results

### Female lungs harbor increased numbers of ILC2s and ILC1s during IAV infection

To determine if the sexual dimorphism in lung ILC2s is maintained during IAV infection, we compared the numbers and phenotype of lung ILCs in 12-14 week old wild-type C57BL/6 female and male mice infected intranasally with a sublethal dose of A/PuertoRico/8/1934 (PR8) virus. We followed weight loss, which peaked at days 7-9 post-infection (p.i.), as a sign of morbidity (Fig. S1A). Females tended to greater weight loss, yet both sexes recovered to near their initial weight by day 12 p.i. Measurement of lung function on day 12 p.i. using a Flexivent instrument showed that females had increased airway resistance, indicative of reduced lung function, suggesting a delay in recovery (Fig. S1B). We used flow cytometry to analyze lung ILCs in independent experiments terminated on days 3, 5, 7,10 and 17 p.i. (Fig. 1; Fig. S1). We defined ILC subsets as lineage-negative (Lin^−^) lymphocytes that expressed CD90, IL7Rα and fate determining transcription factors: T-BET for ILC1s, GATA-3 for ILC2s and RORγt for ILC3s, and we identified NK cells as Lin^−^ CD90^int^ IL7Rα^−^ cells expressing both EOMES and T-BET (Fig. 1A). As in homeostasis, female lungs contained significantly higher numbers of ILC2s compared to males on days 3, 5, 7 and 10 p.i. (Fig. 1B). Relative to naïve mice, numbers of ILC2s were increased at days 5 and 10 p.i., but not at day 7 p.i. when mice show peak morbidity as assessed by weight loss (Fig. 1B, Fig. S1A). By day 17, numbers of ILC2s had returned to levels in naïve mice, with females again harboring significantly higher ILC2 numbers (Fig. S1E). While sex differences in ILC1 numbers in homeostasis were not apparent, female mice showed significantly increased numbers of ILC1s on day 7 p.i. (Fig.1C). Numbers of ILC3s numbers did not differ between sexes in homeostasis and post-infection (Fig. S1C). Notably, males harbored significantly higher numbers of NK cells on days 5 and 7 p.i. (Fig. S1D) consistent with a recent report (29). Thus sex differences in the numbers of ILC2s are retained during IAV infection, with decreased numbers in both sexes on day 7 p.i. relative to day 5 p.i., likely due to reduced levels of GATA3 that make it harder to score cells as ILC2s. Unlike in homeostasis, females have increased ILC1 numbers while males have increased NK cell numbers on day 7 p.i.

**Fig. 1.**
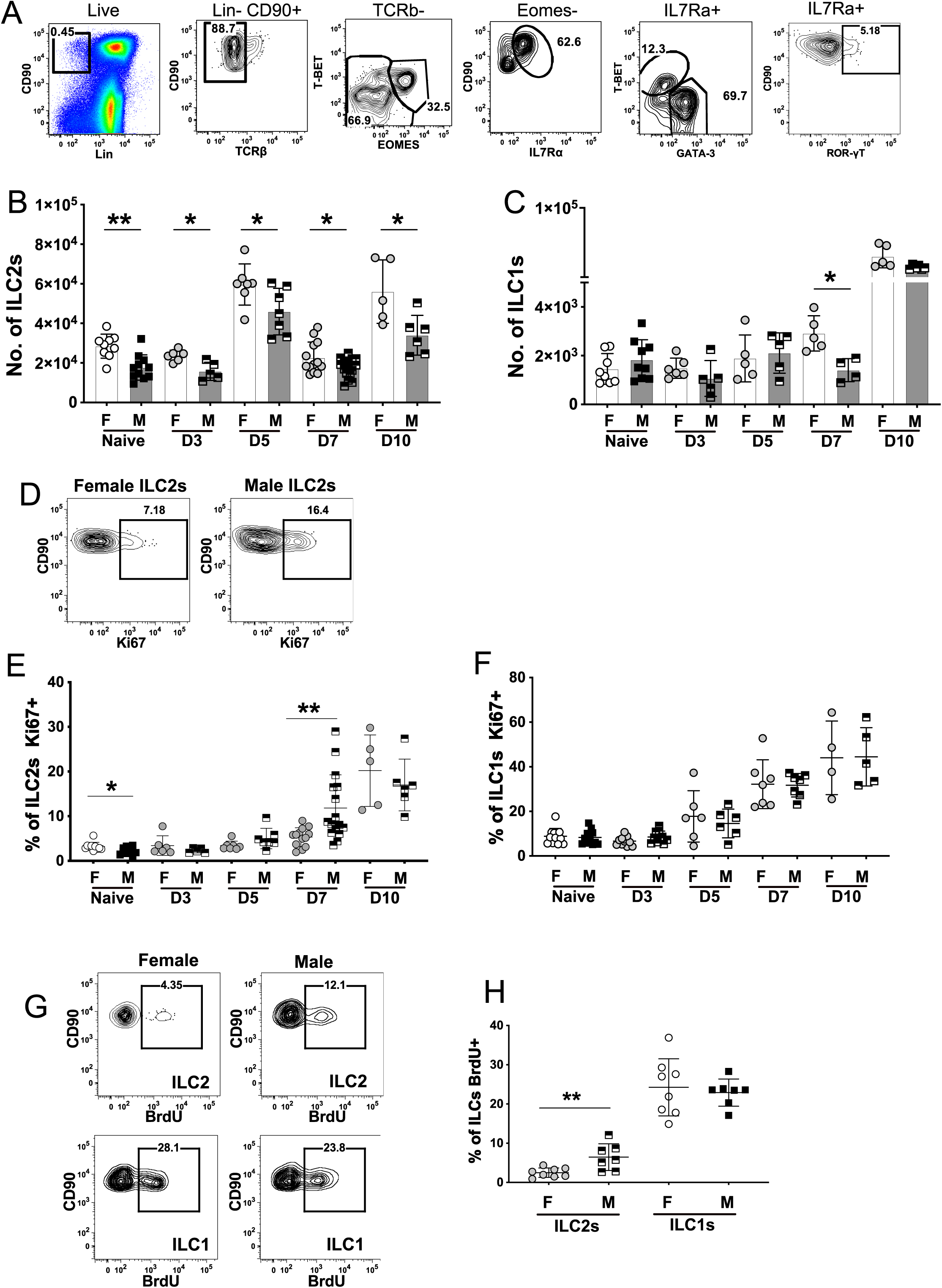
ILC2s in female mice show suppressed proliferation at the peak of influenza virus infection. (**A**) Identification of lung ILCs in a male mouse at day 7 p.i. ILC2s were defined as Lin^−^ CD90^+^TCRβ^−^IL-7Rα^+^ GATA-3^+^, ILC1s were defined as Lin^−^ CD90^+^TCRβ^−^IL-7Rα^+^ T-BET^+^EOMES^−^, ILC3s were defined as Lin^−^ CD90^+^TCRβ^−^IL-7Rα^+^ RORγT^+^, and NK cells were defined as Lin^−^ CD90^+^TCRβ^−^T-BET^+^EOMES^+^. (**B**-**C**) Total numbers of lung ILC2s and ILC1s in naïve and infected female (F) and male (M) mice on the indicated days p.i. (**D**) Identification of Ki67^+^ cells in female and male ILC2s on d.7 p.i. (**E-F**) The fraction of Ki67^+^ ILC2s and ILC1s in naïve and infected female and male mice on the indicated days p.i. (**G**) Identification of BrdU^+^ ILC2s and ILC1s in males and female lungs on d.7 p.i. (**H**) The fraction of BrdU^+^ ILC2s and ILC1s in female and male mice on d.7 p.i. For panels B,C,E,F,H, the data at each time point were compiled from 1-3 independent experiments. Symbols represent individual mice, with the mean and SD indicated. As each time point is an independent experiment, significant differences between males and females at each time point were evaluated using a student t-test for n≥6 and a Mann-Whitney test for n<6.

### ILC2 proliferation is preferentially suppressed in females at the peak of IAV infection

IL-33 released during lung tissue damage induces proliferation of lung ILC2s, yet interferons elicited by IAV restrain ILC2 proliferation (18). Our prior study with a sublethal dose of PR8 virus showed that IFNγ levels peak in the lungs on day 7 p.i. and wane by day 10 p.i. (30). To determine if greater numbers of ILC2s and ILC1s in females after infection correlated with sex differences in ILC proliferation, we measured the percentage of ILCs bearing Ki67, a chromosomal marker associated with progression through the cell cycle. In homeostasis, the percentage of Ki67^+^ ILC2s was significantly higher in female mice, consistent with higher numbers of lung ILC2s in females (Fig. 1E). In contrast, by day 7 p.i., females showed a significantly lower fraction of Ki67^+^ ILC2s compared to males (Fig. 1D,E). By day 10 p.i., sex differences were no longer apparent, and by day 17 p.i., females harbored more Ki67^+^ ILC2s than naïve female mice (Fig. 1E; Fig. S1F). Despite higher numbers of ILC1s in females at later time points p.i., sex differences in the percentage of Ki67^+^ ILC1s were not observed in homeostasis or after infection (Fig. 1F).

We also assayed proliferation of ILC1s and ILC2s during infection using an *in vivo* BrdU incorporation assay. Compared to males, females harbored a significantly lower fraction of BrdU^+^ ILC2s on day 7 p.i. (Fig. 1G,H). In contrast, the frequency of BrdU^+^ ILC1s did not differ between the sexes on day 7 p.i. (Fig. 1G,H). These data show that proliferation of female ILC2s is suppressed relative to that of males at the peak of infection, while ILC1 proliferation is comparable between the sexes.

### In IAV infection, female ILC2s express lower levels of GATA3 and IL-33R/ST2 compared to males

Prior studies showed that levels of GATA3 and IL-33R/ST2 decrease in ILC2s during IAV infection (14). To ascertain sex differences, we measured the relative expression (indicated by mean fluorescence intensity, MFI) of GATA3 and ST2 in ILC2s in infected females and males. Female and male ILC2s did not show differences in the expression of GATA3 in homeostasis, as published (Fig. 2A) (23). On days 5 and 7 p.i., female ILC2s showed significantly lower expression of GATA3 (Fig. 2A,B), but this sex difference was no longer apparent on day 10 p.i. (Fig. 2A). Female ILC2s also showed significantly lower expression of ST2 in homeostasis and on day 7 p.i. (Fig. 2C). Thus, relative to males, female ILC2s have decreased expression of the GATA3 and ST2 molecules associated with ILC2 maintenance and function at the peak of infection (days 5 and 7).

**Fig. 2.**
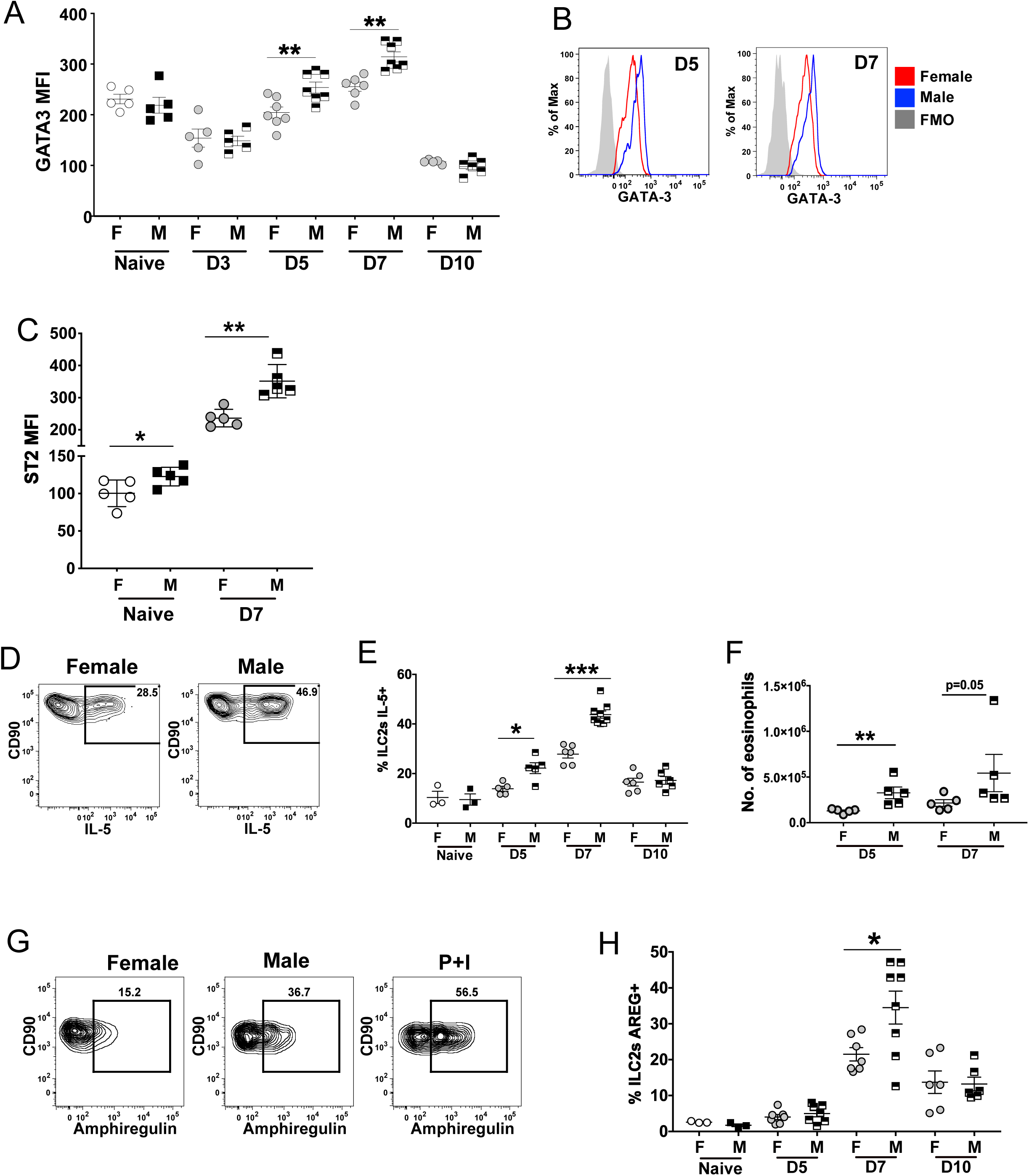
ILC2s in female mice show reduced GATA3 and ST2 expression and a decreased fraction produce IL-5 and AREG. (**A**) GATA3 MFI values in ILC2s of naïve and infected mice on each day. (**B**) Levels of GATA3 expression in male and female lung ILC2s on d.5 and d.7 p.i. relative to an FMO control. (**C**) ST2 levels in male and female lung ILC2s in naïve and d.7 p.i. mice. (**D**) Identification of IL-5 expression in male and male ILC2s on d.7 p.i. without further stimulation *ex vivo*. (**E**) Fraction of IL-5^+^ ILC2s (measured without further stimulation *ex vivo*) on the indicated day p.i. (**F**) Number of eosinophils in male and female lungs at the indicated day p.i. (**G**) Identification of AREG expression in male and male ILC2s on d.7 p.i. without further stimulation *ex vivo*. AREG production by male ILC2s after PMA/Ionomycin (P+I) activation is shown as a positive control. (**H**) Fraction of AREG^+^ ILC2s on the indicated day p.i. Data were compiled from 2-3 independent experiments, and symbols represent individual mice, with the mean and SD indicated. As each time point is an independent experiment, significant differences between males and females at each time point were evaluated using a student t-test for n≥6 and a Mann-Whitney test for n<6.

### A reduced fraction of female ILC2s produce type 2 cytokines during IAV infection

Lung ILC2s activated during IAV infection produce type 2 cytokines including IL-5 and AREG, which promote eosinophil recruitment and tissue repair (11). We hypothesized that the reduced GATA-3 expression in females ILC2s would attenuate their capacity for cytokine production. Indeed, a lower fraction of female ILC2s produced IL-5 on days 5 and 7 p.i. compared to male ILC2s (Fig. 2D,E). Consistent with this, lungs of females contained fewer eosinophils (CD11b^+^SIGLEC-F^+^CD24^+^MHCII^−^) on days 5 and 7 p.i. (Fig. 2F). Production of AREG by ILC2s peaked at day 7 p.i., and compared to males, a lower fraction of female ILC2s produced AREG (Fig. 2G,H). These data show that female ILC2s are less likely to produce type 2 cytokines at the peak of IAV infection, which is consistent with greater suppression of the ILC2 functional program in females.

By day 17 p.i., males and females contained comparable fractions of AREG (∼50%) and IL-5 (∼10%) expressing ILC2s (*not shown*). However, the average number of AREG^+^ ILC2s was 1.8-fold greater in females than in males, and 2.2-fold higher than in naïve mice (Fig. S1G). Thus, once mice have recovered from infection, female ILC2 suppression is alleviated, and a higher number of ILC2s in females continue to produce AREG.

### IAV infected females retain the increased fraction of KLRG1^−^ ILC2s seen in homeostasis

Naïve adult females harbor a prominent KLRG1^−^ subset (∼50-60%) of lung ILC2s that arises after sexual maturity, is negatively regulated by androgens and has been considered immature (23). We hypothesized that IAV infection would decrease the fraction of KLRG1^−^ ILC2s in females, possibly due to activation-induced upregulation of KLRG1 or the preferential proliferation of KLRG1^+^ ILC2s. However, the ratio of KLRG1^−^ and KLRG1^+^ ILC2s remained constant in females and males between days 0 and 10 p.i. (Fig. S2A,B). KLRG1^+^ and KLRG1^−^ ILC2 subsets in females showed no significant differences in proliferation (fraction Ki67^+^ or BrdU^+^) or levels of GATA3 on days 0, 3, 5 and 7 p.i. (Fig. S2C-E). A modestly higher fraction of female KLRG1^+^ ILC2s produced IL-5 compared to KLRG1^−^ ILC2s on day 5 p.i., but this percentage was lower than the fraction of IL-5-producing ILC2s in males that are predominantly KLRG1^+^ (Fig. S2F, 2E). Thus, during IAV infection, females maintain subsets of KLRG1^+^ and KLRG1^−^ lung ILC2s that are both functionally suppressed compared to male ILC2s.

### The preferential female ILC2 suppression is not associated with sex differences in persistence of viral RNA or levels of ILC2 regulating cytokines in the lungs

We next asked if sex differences in viral RNA persistence or the levels of cytokines impacting ILC2 responses correlated with the sex differences in ILC2 function and phenotype observed most notably on day 7 p.i. Viral HA or M1 gene expression in total lung RNA on days 1, 2, 3 or 5 p.i. did not significantly differ by sex (Fig. 3A; Fig. S3A,B). We assessed the levels of cytokine RNAs in total lung cell RNA by qPCR and/or levels of cytokine proteins in the bronchoalveolar lavage fluid (BALF) at early (24-48 hr) and later (days 3-7) time points p.i. Levels of *Il33* RNA at 24-48 hr or on day 7 p.i. did not differ between sexes (Fig. 3B). Levels of *Il12a* and *Il18* RNAs (*not shown*) or levels of IL-18 and IL-1β protein in the BALF on days 5 and 7 p.i. also did not differ by sex (Fig. 3C).

**Fig. 3.**
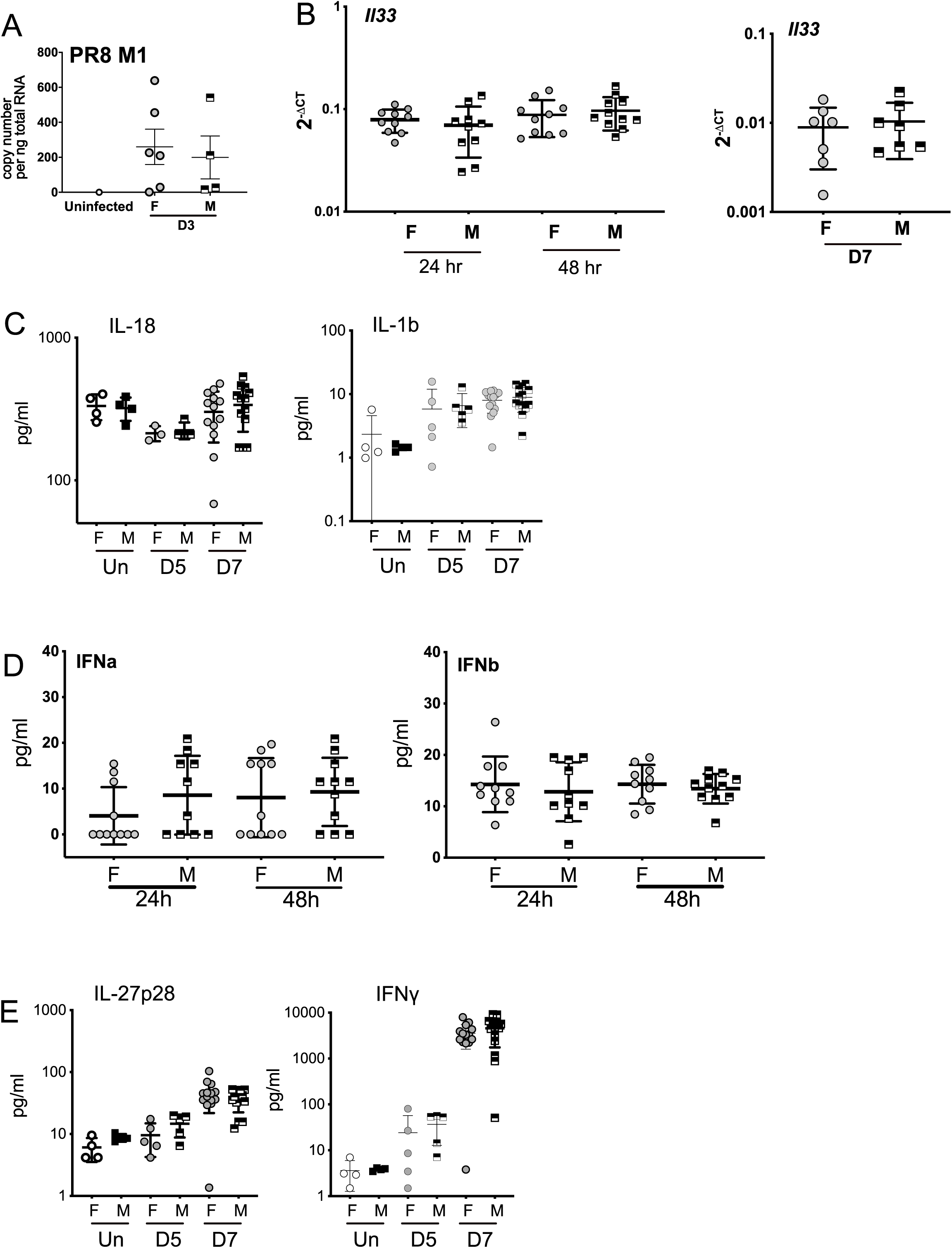
Infected male and female mice do not show significant differences in viral load, IFN or pro-inflammatory cytokines post-infection. (**A**) Copy number of the PR8 M1 RNA in lungs on d.3 p.i. (**B**) Quantification of *Il33* RNA at 24-48 hr and on d.7 p.i. (**C**) Quantification of IL-18 and IL-1β levels in uninfected mice and on d.5-7 p.i. using xMAP assays. (**D**) Quantification of IFNα and IFNβ levels at 24-48 hr p.i. using xMAP assays. (**E**) Quantification of IL-27p28 and IFNγ levels in uninfected mice and on d.5-7 p.i. using xMAP assays. Data were compiled from 2-3 independent experiments, and symbols represent individual mice, with the mean and SD indicated. Significance was evaluated using a one-way ANOVA with a Tukey’s multiple comparison test, n=4-12 per group.

Proliferation and functional responses of ILC2s are restrained in the presence of IFNβ, IFNγ and IL-27 (14–22). The expression of *Ifnb, Ifng* and *Il27* RNAs on days 5 and 7 p.i. did not differ significantly by sex (Fig. S3C). Consistent with this, we did not identify sex differences in BALF levels of IFNα or IFNβ at 24-48 hr p.i. (Fig. 3D) nor in levels of IFNγ or IL-27p28 on days 5 and 7 p.i., (Fig. 3E); IFNα was not detectable on day 7 p.i. We also did not observe significant sex differences in the numbers of CD4^+^ and CD8^+^ T cells producing IFNγ on day 7 p.i. (Fig. S3D) nor the numbers of NP-specific CD4^+^ and CD8^+^ T cells capable of producing IFNγ in an antigen-restimulation assay of lung cells on day 12 p.i. (Fig. S3E). These data suggest that sex differences in ILC2 phenotypes and function on day 7 p.i. are not primarily driven by sex differences in ILC2-suppressing cytokines at the sublethal dose of PR8 virus used. In support of this, we did not find a correlation between BALF IFNγ levels and the fraction of IL-5^+^ ILC2s in either females and males on day 7 p.i. (Fig. S3F).

### Single-cell RNA sequencing identifies sex differences in active and suppressed ILC clusters during IAV infection

To investigate the sex differences in the ILC transcriptome, we performed single-cell RNA sequencing on ∼7800 sorted ILCs (Lin^−^ CD90^+^ IL-7Rα^+^) pooled from females and males on day 7 p.i. Data from two independent experiments were analyzed together (Fig. S4A). Female and male ILCs were identified using antibody-associated hashtags and expression of sex chromosome genes. We clustered all cells using an unbiased graph-based Louvain algorithm and tested gene expression between each cluster to find cluster-specific marker genes via a Wilcoxon rank-sum test (*FindAllMarkers* in *Seurat*). We identified eight different clusters (0–7) with distinct patterns of gene expression (Fig. 4A). Clusters 0, 1, 2 and 4 demonstrated strongly significant (adjusted *p*-value < 1^-176^) differential upregulation of classical ILC2 markers *Gata3* and *Il1rl1* when compared to Clusters 3, 5 and 6 (Fig. 4C) (31). Female and male cells differed in abundance in each cluster (Fig. 4B, S4B).

**Fig. 4.**
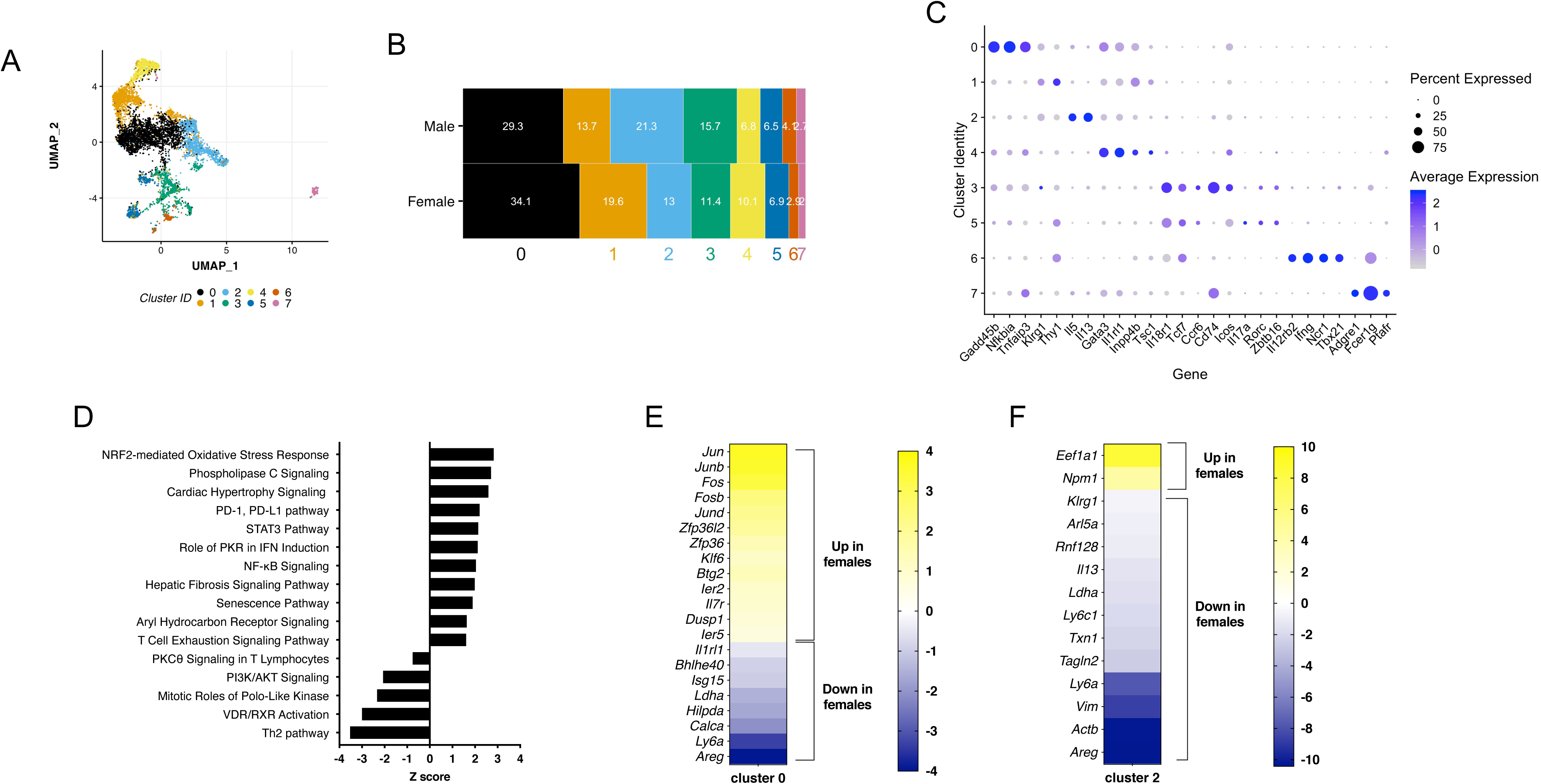
Single cell RNA sequencing identifies sex differences in active and suppressed ILC clusters during IAV infection. Lung ILCs (Lin^−^CD90^+^TCRβ^−^IL-7Rα^+^) were sorted from female and male mice on d.7 p.i. and pooled for scRNA sequencing. Data from 2 independent experiments were put together for subsequent analyses. (**A**) UMAP embedding of PCA projection colored by Louvain clustering. Each cluster retains cells from both experiments. (**B**) Cell type composition by Seurat cluster; numbers indicate the percentage of cells in each cluster in males and females. (**C**) Expression of cluster-specific marker genes. The circle size indicates the percentage of cells within a cluster expressing a gene, while the shading indicates the average expression of the gene within the cluster, scaled across all cells in the population. (**D**) Ingenuity pathway analyses to compare genes in clusters 1 to 2 shows pathways increased in predominantly female cluster 1 (positive Z score) or decreased in cluster 1 (negative Z score) relative to predominantly male cluster 2. (**E**) Sex differences in gene expression within cluster 0 (suppressed ILC2s) identified using the MAST method. Shown is the average difference (log_2_FC). (**F**) Sex differences in gene expression within cluster 2 (activated ILC2s) identified using the MAST method.

Notably, female ILC2s were enriched in clusters 0 and 1 and reduced in cluster 2 relative to male ILC2s (Fig. 4B, S4B). Cluster 0 showed enriched expression of the *Gadd45b*, *Nfkbia* and *Tnfaip3* genes associated with stress and innate immune signaling, and cluster 1 showed enriched expression of *Inpp4b*, a phosphatase known to inhibit IL-5 production, suggesting these clusters represent steady-state or basally activated ILC2s (Fig. 4C). Male cells were enriched in cluster 2 marked by genes involved in canonical ILC2 function such as *Il5, Il13* and *Areg* (Fig. 4C). We then mapped a previously published scRNA-seq dataset acquired from a pool of female and male lung ILC2s in homeostasis (32) to our defined clusters (Fig. S4D). This analysis showed that the vast majority of cells from the published dataset fell into our defined clusters 0 and 1. This comparison suggests that our clusters 0 and 1 might best represent a pool of pre-infection ILC2s present in the lung. However, Ingenuity pathway analyses revealed that genes increased in the female-biased cluster 1 relative to the male-biased cluster 2 mapped to pathways involved in NRF2-mediated oxidative stress response, IFN induction, senescence and exhaustion (Fig. 4D). Genes decreased in cluster 1 relative to cluster 2 mapped to PKCθ or PI3K/AKT signaling and Th2 pathways (Fig. 4D). These unbiased transcriptome data reinforce our cellular assay findings that female ILC2s either remained in a state typical of homeostasis or were more functionally suppressed than male ILC2s.

Clusters 3, 5 and 6 showed strongly significant (adjusted *p*-value ≤ 1^-159^) differential expression of genes typical of α-lymphoid precursors, ILC1s and ILC3s, including *Il18r, Zbtb16* and *Tcf7* (Fig. 4C) (31). Cluster 3 was typified by expression of *Ccr6*, *Icos*, *Il18r1*, *Cd74,* and MHCII genes, yet lacked *Ncr1*, characteristics of activated, antigen-presenting ILC3s (33). Cluster 5 showed the highest expression of canonical ILC3 genes *Il17a*, *Rorc* and *Il18r*. Cluster 6 showed high expression of canonical ILC1 and NK cell genes *Il12rb2*, *Ifng*, *Ncr1, Eomes* and *Tbx21*. Cluster 7, distinguished by *Adgre1*, *Fcer1g*, *Il1b* and *Ptafr*, likely represents a contaminating myeloid population.

To identify sex differences in gene expression in each cluster, within-cluster female-male differential gene expression testing was performed using the *MAST* method. Within cluster 0 (Fig. 4E), genes increased in females include a group of conserved coregulated early response genes (*Jun, Junb, Jund, Fos, Fosb, Ier2, Btg2, Dusp1*) also associated with expression of *Zfp36* and *Zfp36l2* (encoding tristetraprolin), RNA-binding molecules that promote mRNA degradation of proteins involved in inflammation and proliferation, and are important to the attenuation of T cell activation (34–36). Notably, both *Zfp36* and *Zfp36l2* transcripts were significantly elevated in female compared to male cells in cluster 0. In contrast, genes involved in proliferation (*Ldha, Isg15*) and ILC2 function (*Areg, Calca, Bhlhe40, Il1rl1*) were decreased in female relative to male cells in cluster 0 (37–40). Within cluster 2 (Fig. 4F), genes decreased in female compared to male cells include those involved in ILC2 function and downstream of GATA3 (*Areg, Il13, Rnf128*), proliferation (*Ldha, Txn1*) and cytoskeletal rearrangement (*Actb, Vim, Tagln2*) (41). Taken together, these data show that female ILC2s that fall into cluster 2 marking activated ILC2s show decreased expression of genes associated with ILC2 function or proliferation compared to male cells in cluster 2, while male ILC2s that fall into cluster 0 marking resting or suppressed ILC2s show increased expression of genes in pathways involving ILC2 function and proliferation compared to female cells. Selected genes (*Ly6a, Areg, Ldha*) are decreased in females in both clusters, suggesting sex-specific regulation of genes important to ILC2 biology.

### Female lung resident ILC2s show intrinsic differences in IFNGR expression and STAT1 signaling in homeostasis and infection

Having established notable sex differences in ILC2 activation phenotypes during IAV infection, we next tested the hypothesis that the increased ILC2 suppression in females stems from ILC2-intrinsic sex differences in response to IFN. We first assessed IFN receptor expression in lung ILC2s in naïve and infected mice using flow cytometry. Female ILC2s expressed significantly higher levels of IFNGR (judged by MFI) than male ILC2s in homeostasis (Fig. 5A,B). In contrast, ILC1 expression of IFNGR did not differ between the sexes (Fig. 5C). On day 7 p.i., levels of IFNGR were significantly higher in female compared to male ILC2s, while ILC1s did not show sex differences in IFNGR levels (Fig. 5B,C). IFNGR expression did not differ between KLRG1^+^ and KLRG1^−^ ILC2s in naive females (Fig. S2G). We did not detect expression of the type I IFN receptor in ILC2s using flow cytometry. The differences in IFNGR expression in homeostasis suggested that females and males will have cell intrinsic differences in the magnitude of IFNGR signaling.

**Fig. 5.**
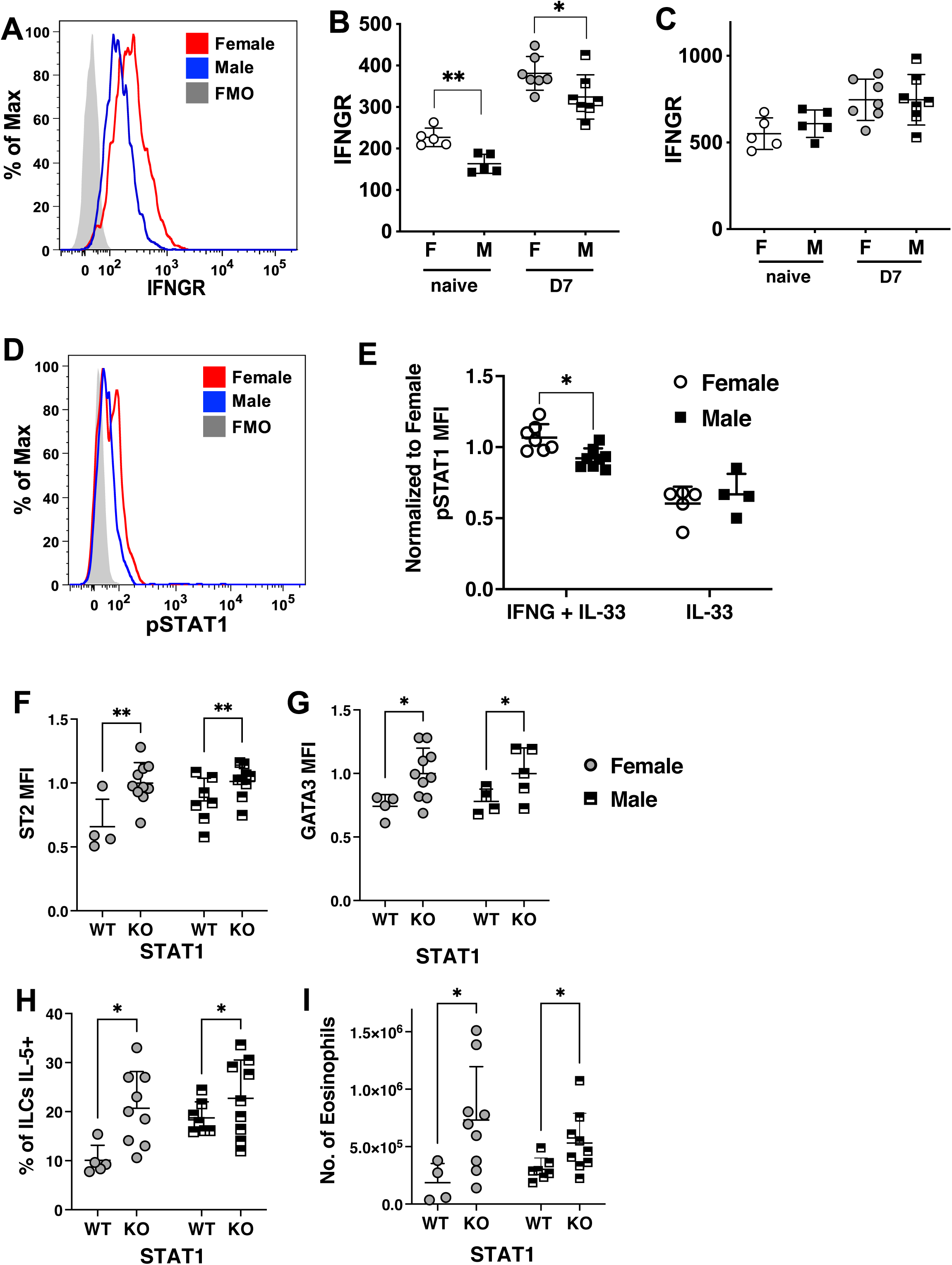
Female ILC2s show intrinsic differences in IFNGR expression and STAT1 signaling in homeostasis and infection. (**A**) Expression of IFNGR on female and male lung ILC2s in naive mice relative to the FMO control. (**B-C**) Compiled expression (MFI) of IFNGR on female and male lung ILC2s and ILC1s in naïve or infected (d.7 p.i.) mice. Symbols represent individual mice, with the mean and SD indicated. As each time point is an independent experiment, significant differences between males and females at each time point were evaluated using a Mann-Whitney test. (**D**) Intracellular staining for pSTAT1 in female or male ILC2s stimulated in vitro with IFNγ, relative to an FMO control. (**E**) pSTAT1 MFI in ILC2s normalized to the average value in IFNγ stimulated female cells. Data were compiled from 2 independent experiments and symbols represent individual mice, with the mean and SD indicated. Significance was evaluated using a mixed effects model 2 way ANOVA comparing sex and treatment followed by a Tukey’s multiple comparison test. (**F-I**) ILC2s in lungs of female and male *IL-7R-cre-Stat1^+/+^* (WT) and *IL-7R-cre-Stat1^f/f^* (KO) mice were analyzed on d.7 p.i. (**F**) ST2 MFI normalized to the average value in KO ILC2s in females. (**G**) GATA3 MFI normalized to the average value in KO ILC2s in females. (**H**) Fraction of ILC2s expressing IL-5. (**I**) Number of eosinophils. Significance in panels F-I was evaluated using a two way ANOVA comparing sex and genotype with a Tukey’s multiple comparison test.

IFNγ binding to IFNGR induces phosphorylation of the transcription factor STAT1, which forms a homodimer that translocates to the nucleus and binds to IFNγ activated sites in DNA (42). To determine if ILC2s show sex differences in STAT1 activation, we measured expression of phosphorylated (Y701) STAT1 (p-STAT1). Naïve lung ILC2s of each sex were sorted and stimulated with vehicle or IFNγ for 15 min. in the presence of IL-33 and IL-7. Notably, IFNγ stimulation led to a subset of ILC2s in females showing significantly higher levels of p-STAT1 in female ILC2s (Fig. 5D,E).

To determine if STAT1 is required for female ILC2 suppression during IAV infection, we generated mice bearing either conditional or wild-type alleles of *Stat1* bred to IL-7R-cre (“STAT1-deficient”, IL-7R-cre-Stat1^fl/fl^ or “wild-type, WT” IL-7R-cre-Stat1^+/+^) (Fig. 5F-I). Compared to WT female ILC2s, STAT1-deficient female ILC2s showed higher levels of ST2 and GATA3 that were comparable to levels in WT male ILC2s (Fig. 5F,G), indicating that the absence of STAT1 signaling prevented suppression of female ILC2s. Similarly, the fraction of IL-5^+^ ILC2s was comparable in STAT1-deficient female and WT male mice (Fig. 5H), which correlated with an increased number of eosinophils (Fig. 5J). STAT1 deficiency in male ILC2s also increased their activation to a modest degree (Fig. 5G-I). Taken together, these data support the hypothesis that female ILC2s are subject to greater suppression during IAV infection through elevated IFNGR signaling mediated via STAT1.

### Androgens regulate the sex differences in numbers, phenotype and cytokine production of ILC2s during IAV infection

Androgens negatively regulate lung ILC2 numbers, phenotype and functional responses in homeostasis and type 2 inflammation models. To test the hypothesis that endogenous androgens in males regulate sex differences in ILC2 responses during IAV infection, we performed orchiectomy (castration, Cx) of young (4-5 weeks old) males to remove the major source of androgens prior to sexual maturity. Males were infected at 14-16 weeks of age. Notably, Cx males showed a dramatic increase in lung ILC2 numbers compared to intact males on day 7 p.i. (Fig. 6A). In Cx males, the number of KLRG1^−^ ILC2s was comparable to the number in females on day 7 p.i., indicating that androgens in males inhibit the accumulation of the KLRG1^−^ ILC2 subset that is prominent in female lungs (Fig. S5A,B). Cx and intact males showed no differences in the numbers of ILC1s or NK cells on day 7 p.i. (Fig. S5C,D), suggesting that androgens do not directly regulate the sex differences in their numbers.

**Fig. 6.**
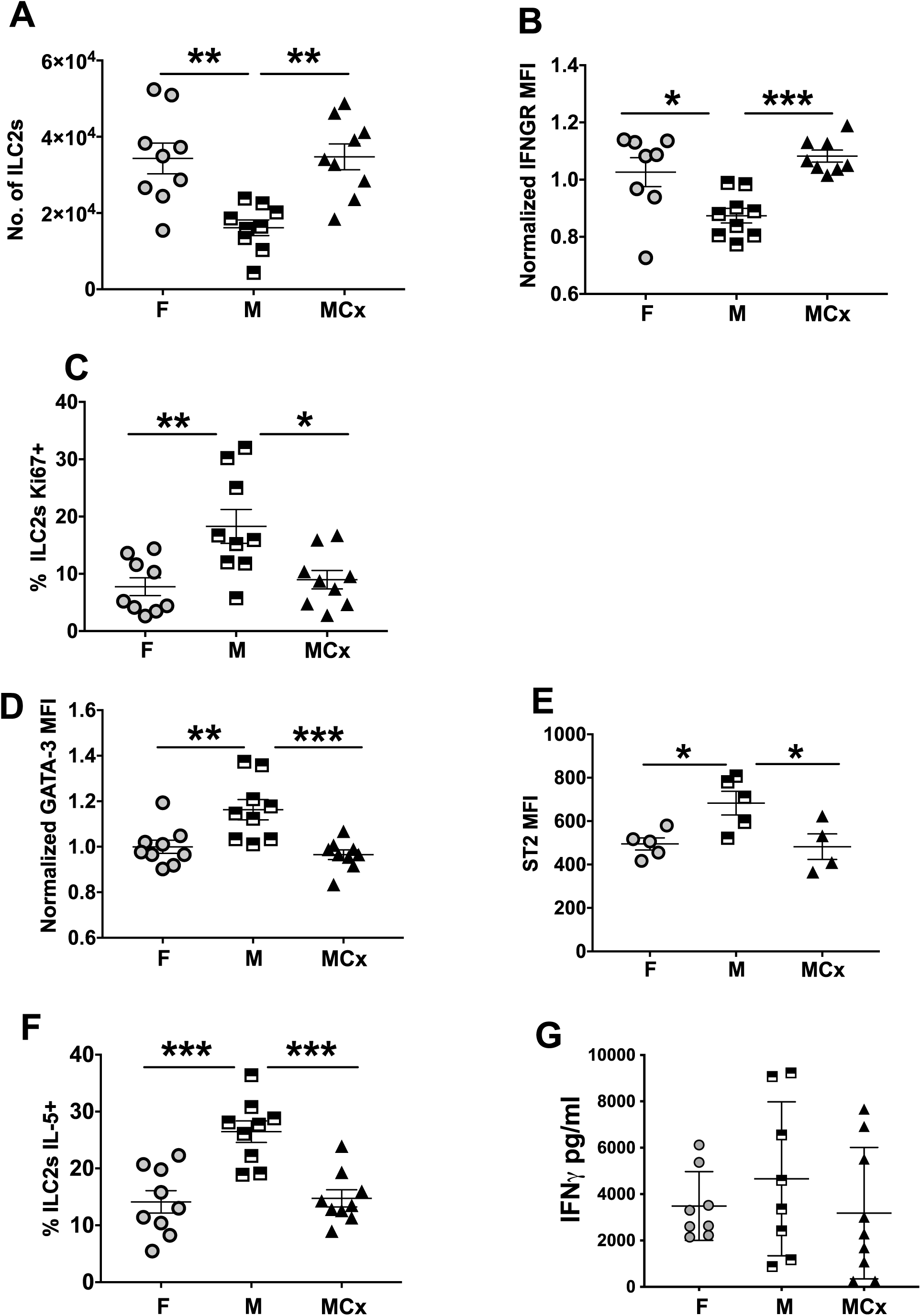
Male ILC2s adopt the phenotype of female ILC2s in orchiectomized mice. Male mice were orchiectomized (MCx) at 4 weeks of age and infected with IAV at 14 weeks of age. Lung ILC2s were analyzed at d.7 p.i. (**A**) Total number of ILC2s in the lung in females (F), males (M) and orchiectomized males (MCx). (**B**) MFI of IFNGR on ILC2s (normalized to female average). (**C**) Fraction of ILC2s that were Ki67^+^. (**D**) MFI of GATA3 on ILC2s (normalized to female average). (**E**) MFI of ST2 on ILC2s. (**F**) Fraction of ILC2s that were IL-5^+^. (**G**) IFNγ levels in BAL. Data are compiled from 1-2 independent experiments, and symbols represent individual mice. Significance was evaluated using a one way ANOVA with a Tukey’s multiple comparison test.

Androgens also regulated the sex differences in suppression of lung ILC2s on day 7 p.i. Notably, we observed increased expression of IFNGR in ILC2s of Cx males compared to intact males, with levels comparable to IFNGR on female ILC2s (Fig. 6B). ILC2s in Cx males showed a significantly lower fraction of Ki67^+^ cells and lower levels of GATA-3 and ST2 compared to ILC2s in intact males (Fig. 6C-E). The fraction of IL-5-producing ILC2s also was significantly reduced in Cx males, to a level comparable to females (Fig. 6F). This correlated with a trend to reduced numbers of eosinophils and a decreased frequency of AREG^+^ ILC2s in Cx males compared to intact males (Fig. S5E,F). Levels of IFNγ (Fig. 6G) and IL-18 (Fig. S5G) in BAL fluid did not differ between intact and Cx males and females. These data show that androgens preserve canonical ILC2 responses during IAV infection, and this correlates with reduced expression of IFNGR in ILC2s.

### Androgen receptors regulate the sex differences in ILC2 numbers and phenotypes

Lung ILC2s express high levels of androgen receptor (*Ar*) but very low levels of estrogen receptor (*Esr1* and *Esr2)* RNAs (9). Here we used intracellular staining with an anti-AR mAb to show that lung ILC2s preferentially express AR protein, with significantly higher levels in male ILC2s (Fig. 7A,B). In contrast, ILC1s and B and T lymphocytes in the lungs did not show detectable levels of AR protein (Fig. 7B).

**Fig. 7.**
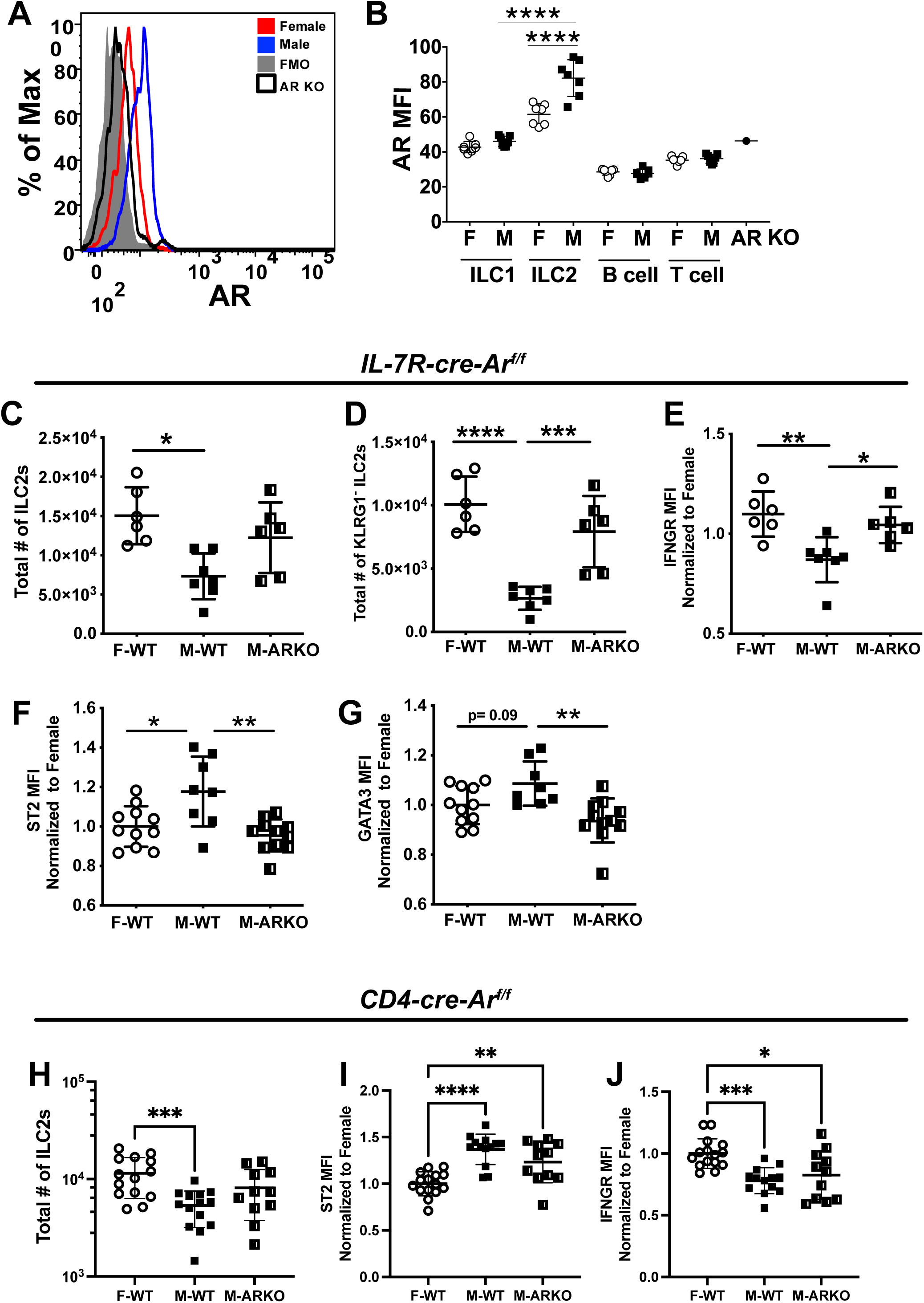
ILC2s express high levels of AR, and male ILC2s lacking AR adopt phenotypes of female ILC2s. (**A**) Staining with anti-AR mAb in female and male lung ILC2s in homeostasis, relative to an FMO control and the background binding of the mAb to ILC2s in *IL-7R-cre-Ar^fl/fl^* mice (AR KO). (**B**) Compilation of AR MFI values on lung lymphocyte subsets in females (F) and males (M) in homeostasis. (**C-E**) Lung ILC2s were assessed during homeostasis in female, male WT and *IL-7R-cre-Ar^fl/fl^*mice (AR KO): (**C**) Number of lung ILC2s. (**D**) Number of KLRG1^−^ ILC2s. (**E**) IFNGR MFI. (**F-G**) Lung ILC2s were assessed on d.7 p.i. with IAV in female, male WT and *IL-7R-cre-Ar^fl/fl^*mice (AR KO): (**F**) ST2 MFI. (**G**) GATA3 MFI. (**H-J**) Lung ILC2s were assessed on d.7 p.i. with IAV in female, male WT and *CD4-cre-Ar^fl/fl^* mice (AR KO): (**H**) Number of lung ILC2s. (**I**) ST2 MFI. (**J**) IFNGR MFI. Data are compiled from 2-3 independent experiments, and symbols represent individual mice, with mean and SD indicated. Significance was evaluated using a one way ANOVA with a Tukey’s multiple comparison test.

To test the hypothesis that loss of lymphocyte-restricted AR activity accounts for one or more of the effects of orchiectomy on male ILC2 phenotypes, we bred mice bearing wild-type or conditional alleles of *Ar* and IL-7R-cre (“AR-deficient” IL-7R-cre-Ar^fl/fl^ or “wild-type, WT” IL-7R-cre-Ar^+/+^). In homeostasis, ILC2 numbers in male AR-deficient mice were comparable to those in female WT mice, which was primarily due to significantly increased numbers of KLRG1^−^ ILC2s (Fig. 7C,D). ILC2s in naïve male AR-deficient mice contained IFNGR and GATA3 at higher levels than in WT male ILC2s and comparable to WT female ILC2s (Fig. 7E), suggesting that AR activity decreases expression of IFNGR and GATA3 in naïve ILC2s. In IAV infected mice on day 7 p.i., AR-deficient ILC2s in males showed levels of reduced ST2 and GATA3 levels comparable to that of female ILC2s (Fig. 7F,G).

IL-7R-Cre drives deletion of *Ar* in all lymphocytes starting in the common lymphoid progenitor. Therefore, to determine if AR deficiency in T cells, but not ILCs, might alter ILC2 phenotypes, we bred mice bearing CD4-Cre to delete *Ar* in CD4^+^ and CD8^+^ T cells. In IAV infected mice on day 7 p.i., numbers of ILC2s were comparable in male CD4-cre-Ar^fl/fl^ and CD4-cre-Ar^+/+^ lungs (Fig. 7H), indicating that AR deficiency in T cells did not alter the sex difference in ILC2 numbers. Similarly the level of expression of ST2 and IFNGR was comparable on ILC2s in male CD4-cre-Ar^fl/fl^ and CD4-cre-Ar^+/+^ lungs and distinctly different than levels on female ILC2s (Fig. 7I-J). Taken together, these data obtained with mice bearing deletions of *Ar* driven by IL-7R-Cre or CD4-Cre support our hypothesis that ILC2-intrinsic AR protects male ILC2s from suppression during IAV infection.

## Discussion

In IAV infection, lung-resident ILC2s generate type 2 responses that promote viral clearance and repair of injured lung tissue (12,43,44), yet they also may be suppressed or show plasticity in the IFN-centered type 1 inflammatory environment (14–22). Herein, we have demonstrated a sex difference in the extent of ILC2 suppression at the peak of morbidity in a sublethal IAV infection of mice. Compared to those in males, ILC2s in females showed attenuated proliferation, decreased propensity for IL-5 and AREG production, and reduced expression of GATA3 and IL-33R, indicating greater suppression of canonical ILC2 function in females. We also provided evidence that the AR and IFNGR-STAT1 signaling pathways regulate these ILC2-intrinsic sex differences during influenza virus infection.

Our data show that sex biased ILC2 suppression is regulated by androgens and AR activity. Consistent with reports of high *Ar* RNA expression, we found that ILC2s contain high levels of AR protein relative to other lymphocytes in the lung during homeostasis. Our data showing sex differences in ILC2 proliferation during infection are supported by prior reports that AR expression reduces the proliferation of ILC2s (6). We showed that all lymphocyte-restricted (IL-7R-Cre), but not T cell-restricted (CD4-Cre), AR deficiency led to increased levels of IFNGR on male ILC2s, suggesting attenuation of IFNGR expression by AR activity in males. Indeed, depletion of androgens by orchiectomy abolished the sex differences in multiple parameters including IFNGR expression, leading to ILC2 suppression in orchiectomized males that is comparable to females post-infection. ARs are ligand-dependent transcription factors that mediate long-range chromatin interactions by partnering with histone-modifying coactivators and corepressors at gene regulatory elements, thus promoting epigenetic changes and modulating transcription (26). Notably, identification of AR target genes using anti-AR chromatin immunoprecipitation (ChIP) in human prostate cancer cells revealed that AR binds to genes within the IFNGR signaling pathway (*STAT1*, *IRF1*) as well as to genes involved in ILC2 development, maintenance or function (*RORA, ETS1, NFIL3, TCF7, ZBTB16*) (45,46). These findings suggest that sex differences in ILC2 function are imposed by AR activity in males. More broadly, testosterone and male sex have been reported to reduce morbidity and promote recovery of mice infected with moderate doses of influenza virus (5,47–50). Our data show that preservation of pulmonary ILC2 function by testosterone might contribute to the reduced morbidity in males.

Our experiments also support the hypothesis that the biased suppression of female ILC2s is due to increased IFNGR expression and signaling. In homeostasis, we noted sex differences in cytokine receptors with female ILC2s showing higher surface levels of IFNGR and male ILC2s displaying higher surface levels of IL33R/ST2. Naïve female ILC2s showed greater STAT1 phosphorylation upon stimulation by IFNγ *in vitro*, and lymphocyte-restricted STAT1 deficiency reversed suppression of female ILC2s during *in vivo* infection. These findings are consistent with prior reports that suppression of ILC2s during respiratory virus infection is mediated via IFNGR or IFNAR signaling, which signal via STAT1 (16,17,20). As described above, AR has been linked to the STAT1 pathway, raising the possibility that AR activity attenuates transcription of genes in the IFNGR signaling pathway in ILC2s. Consistent with increased ILC2 suppression in IAV infected females, scRNA sequencing showed that female ILC2s in infected lungs preferentially grouped into clusters enriched in pathways involving oxidative stress response, IFN induction, senescence and exhaustion, while male ILC2s preferentially grouped into clusters enriched in the PKCθ or PI3K/AKT signaling and Th2 pathways.

In contrast to the marked suppression of female ILC2 functional responses observed during IAV infection, others observed preferential ILC2 activation in females challenged intranasally with allergens to induce allergic airway inflammation [reviewed in (51)]. In the House Dust Mite extract induced type 2 model, we also found higher numbers of ILC2s that were Ki67^+^ in females compared to males (Fig. S6A,B). A greater fraction of ILC2s in females were IL-5^+^ (Fig. S6C), consistent with a higher number of eosinophils in female lungs (Fig. S6D). The heightened function of female ILC2s in a type 2 model is in sharp contrast to preferential suppression of female ILC2s during type 1 inflammation that we report here.

Sex differences in the functional responses of multiple immune cell types likely contribute to observed sex disparate responses or outcomes in viral infection (52). Our findings raise the question of whether sex differences in suppression of canonical ILC2 function contribute to these documented sex differences during influenza infection or to differences in regeneration of damaged tissue once infection is resolved. Upon a sublethal infection, we found that increased ILC2 suppression and ILC1 numbers in females at the peak of infection correlated with a persistent increase in airway resistance indicative of reduced lung function and a trend to greater weight loss, but we detected little sex difference in viral load or persistence, consistent with earlier reports (4,5). Increased conversion of ILC2s into ILC1-like cells producing IFNγ in preferentially in females may contribute to the level of type 1 inflammation, thus increasing morbidity in influenza infection (14). Indeed, a prior report showed that ILC2s promote survival when IFNγ is neutralized during IAV infection, suggesting canonical type 2 ILC2 functional responses enhance recovery (19).

We have not yet studied sex differences in tissue repair once virus is cleared, but we expect that ILC2 functional responses will regulate this phase. The repair of the epithelium, involving matrix deposition and epithelial cell proliferation and differentiation, is crucial for recovery from infection (44) and is promoted by IL-5 and AREG, both of which are produced by ILC2s during influenza infection (13,53). IL-5 recruits eosinophils that promote anti-viral immunity and lung tissue regeneration (53,54). AREG signals via EGF receptor on epithelial and stem cells leading to their proliferation and differentiation, thus promoting restoration of tissue integrity after viral clearance (13). An early study in T cell deficient mice showed that depletion of ILC2s led to the loss of airway epithelial integrity, diminished lung function and impaired airway remodeling, and the administration of recombinant AREG to these ILC2 and T cell deficient mice improved lung function and airway epithelial integrity (12). Similarly, the absence of ILC2s in *IL7R-cre-Rora^−/–^* mice led to decreased survival during a lethal infection (19). However, global IFNAR deficiency led to increased numbers of ILC2s and type 2 pathology during influenza virus infection, suggesting that ILC2 suppression by IFN might be beneficial for attenuation of pulmonary fibrosis after infection (17). These studies suggest that ILC2s contribute significantly to tissue repair, and that sex differences in canonical ILC2 function resulting in quantitative differences in IL-5 or AREG might impact recovery from infection. A nonredundant role for ILC2 products during infection remains to be established however, as studies show that lung epithelial cells and T regulatory cells also produce AREG after influenza infection (47,55). AREG production in respiratory epithelial cells during influenza infection was increased in males, although AREG and testosterone were determined to have independent effects in protection of males from severe disease (47).

Epidemiological studies show that morbidity rates and hospitalization during influenza virus pandemics were higher in women during their reproductive years (2). In contrast, elderly men have shown greater morbidity and mortality during the COVID-19 pandemic and pandemic influenza episodes (56,57). While the mechanisms underlying these biological sex differences are surely multifactorial and likely include regulation of immune and non-immune cells by sex steroid receptors, our data underscore the compelling need to understand sex differences in baseline immune parameters as well as immune responses upon respiratory virus infection.

## Materials and Methods

### Mice

Female and male C57BL/6J mice were purchased from Jackson Labs (JAX) and used within 2-4 weeks or bred in house. Mice bearing a conditional allele of *Ar* [B6.129S1-Ar<tm2.1Reb>/J] were purchased from JAX and bred to mice bearing IL-7Rα promoter-driven Cre recombinase [kindly provided by Dr. H.R. Rodewald] (58) or to mice bearing CD4 promoter-driven Cre recombinase [Tg(Cd4-cre)1Cwi/BfluJ] (59). Mice bearing a conditional allele of *Stat1* [kindly provided by Dr. L. Olson] (60) were bred to IL-7Rα-Cre mice. Mice were maintained as IL-7Rα-Cre heterozygotes, and control mice contained IL-7Rα-cre and wild-type alleles of *Ar* and *Stat1*. Age-matched females and males between 12-16 weeks of age were compared, with mice taken from the same breeding colony or purchased together. Orchiectomy (castration) was performed on male C57BL/6J mice 4-5 weeks after birth, and the mice were used 8-12 weeks after surgery. All experiments were approved by the OMRF Institutional Animal Care and Use Committee.

### Influenza virus infection

IAV was grown in embryonated chicken eggs, and viral end point titer (egg infectious units, EIU) was determined using allantoic fluid in a hemagglutination assay in the lab of Dr. Gillian Air (University of Oklahoma Health Sciences Center). HA genes were sequenced to confirm identity of virus stocks as described previously (30). Mice were infected intranasally with 30 μl PBS containing 3×10^2^ EIU of mouse adapted A/Puerto Rico/8/1934 (PR8, H1N1) virus after they were anesthetized with ketamine and xylazine. Mice were weighed daily to track morbidity during the infection.

### Tissue Preparation

Lungs were perfused with PBS + 1 mM EDTA, minced and incubated in 100μg/ml of Liberase TM (Roche) and 100μg/ml of DNase I (Roche) at 37°C for 45-60 min, and red blood cells were lysed with ACK buffer (BD Biosciences). A single cell suspension was prepared by passing the tissue through a 70μm filter.

### Administration of BrdU

Mice were infected intranasally with PR8 virus and administered BrdU (0.8 mg per mouse) by intraperitoneal injection at day 4, 5 and 6 p.i. Incorporation of BrdU was detected in lung cells of infected mice at day 7 p.i. by flow cytometry using the manufacturer’s protocol (BD Biosciences).

### Flow cytometry

Complete flow cytometry gating strategies for lung ILCs are shown in Fig. 1 and S1. Dead cells were identified with a fixable Zombie Aqua stain (Biolegend). Surface staining was performed by incubating cells with mAbs on ice for 15 min after 5 min of anti-CD16/32 treatment. The Foxp3/Transcription Factor Staining Buffer Set (ThermoFisher Scientific) for intranuclear staining and the Cytoperm/Cytofix ^TM^ Solution Kit (BD Biosciences) for intracellular staining were used according to manufacturer instructions. Lung ILCs were identified after gating on the singlet, live population and excluding lineage-positive cells using mAbs specific for the lineage markers-CD3 (145-2C11), CD5 (53-1.3), TCRβ (H57-597), B220 (RA3-6B2), CD11b (M1/70), CD11c (N418), TER119 (TER-119), Gr-1 (RB6-8C5), CD19 (MB19-1). Combinations of fluorochrome-conjugated mAbs specific for CD90.2 (30-H12), NK1.1 (PK136), IL7Rα (A7R34), ST2 (RMST2-2), CD25 (PC61.5), KLRG1 (2F1/KLRG1), CD218a (IL-18Rα-Clone P3TUNYA), CD119 (IFNGR-Clone GR20), EOMES (Dan11mag), GATA3 (L50-823), T-BET (4B10), RORγT (Q31-378), BrdU (B44), Ki-67 (B56), IL-5 (TRFK5), AR (EPR1535(2)) and Amphiregulin (BAF989) were used to identify and characterize NK cells and ILCs. Eosinophils were identified using CD11c (N418), SIGLEC-F (E50-2440), CD24 (M1/69), MHCII (M5/114.15.2), and CD11b (M1/70). T cells were identified with mAbs specific for CD3 (145-2C11), CD4 (GK1.5), CD8α (53-6.7), IFNγ (XMG1.2), TNFα (MP6-XT22) and GZMB (GB11). All mAbs were purchased from BD Biosciences, Biolegend, Tonbo Biosciences, Thermofisher Scientific, Abcam and R&D Systems. Samples were acquired on an LSRII instrument containing 4 lasers and analyzed using FlowJo software (Ver. 9.8.5).

### Ex vivo detection of cytokine production

For detection of cytokine production by ILC2s, lung lymphocytes were enriched with 40% Percoll (GE Healthcare Life Sciences) and then incubated in complete medium (RPMI+10% FCS) without further stimulation for 5 hr in presence of monensin (5 μg/ml, BD Biosciences). Buffers optimized for intracellular cytokine staining or intranuclear staining were used to identify IL-5 and AREG production by GATA3^+^ ILC2s. As a positive control, lung cells were incubated with PMA (30 ng/ml) and ionomycin (500 ng/ml) for 5 hr, and monensin added for the last 2.5 hr. T cell stimulation experiments were performed as described (30). Briefly, 2-4×10^6^ lung cells were incubated with the H-2D^b^-binding NP_366-374_ peptide (2 μg/ml) (MBLI) and the I-A^b^-binding NP_311-325_ peptide (2 μg/ml) (Bio-Synthesis Inc) and brefeldin A (5 μg/ml) (BD Biosciences) in RPMI+5% FCS for 5 hr. Surface and intracellular staining identified IFNγ, TNFα or GZMB producing CD4^+^ and CD8^+^ T cells.

### Cell sorting

Lung lymphocytes were enriched with a 40% Percoll gradient after LiberaseTM and DNase I digestion of the total lung. Cells were stained with mAbs to lineage markers-CD3, CD5, TCRβ, B220, CD11b, CD11c, TER119, Gr-1, CD19- and ILC2-specific markers. After gating on the Lin^−^ cells, ILC2s were sorted as CD90^+^IL7Rα^+^ST2^+^CD25^+^ cells into PBS+20% FCS using a BD FACS Aria II instrument.

### Detection of phosphorylated STAT1 in ILC2s

ILC2s were sorted from naïve CD45.2^+^ females and males and stimulated with IFNγ (1 ng/ml) or vehicle at 37°C for 15 min, in the presence of IL-7 (10 ng/ml) and IL-33 (2 ng/ml). After stimulation, cells were immediately fixed in BD Phosflow™ Fix Buffer I following the manufacturer’s protocol. Fixed CD45.1^+^ splenocytes stained with anti-CD45.1 mAb were added as carrier cells. Fixed cells were permeabilized using BD Perm Phosflow™ Buffer III and incubated for 30 min at room temperature with mAbs specific to STAT1 (pY701-Clone 4a) and GATA-3 (L50-823) (BD Biosciences) and analyzed by flow cytometry.

### Isolation of RNA and real-time quantitative PCR

Lung tissue was lysed with Trizol reagent (Ambion), and RNA was isolated with a Trizol-RNeasy mini kit (QIAGEN) hybrid protocol. cDNA was synthesized using iScript™ gDNA Clear cDNA Synthesis kit (BIO-RAD) as described (30). Quantitative real-time RT-PCR (qPCR) of genes was performed on an ABI 7900HT instrument. Expression of tested cytokine genes relative to *Gapdh* or *Eef2* expression was determined using the 2^−ΔCt^ method (30). To quantify viral load, qPCR of the PR8 *HA* gene in lung samples was performed, and expression of the *HA* gene relative to *Eef2* expression was determined using the 2^−ΔCt^ method. PR8-M1 (Matrix protein 1) copy number within lung samples was determined using qPCR. A 469bp region of the PR8 M1 protein was cloned into the plasmid pMiniT 2.0, and this construct was used to generate a standard curve to calculate the amount of viral DNA in samples. The primers are listed in Table S1.

### Cytokine assays

Bronchoalveolar lavage (BAL) fluid was collected by washing the lung cavity with PBS three times using an intratracheal catheter (Tom cat 3.5 Fr x5.5”, Santa Cruz Animal Health). Cytokines in the BAL fluid were measured using xMAP multiplex assays in the OMRF Serum Analyte and Biomarker core facility. ProcartaPlex kits (ThermoFisher Scientific) were used according to the manufacturer’s instructions.

### FlexiVent measure of lung function

Naïve and infected mice were anesthetized with ketamine/xylazine and subjected to a tracheotomy. Cannulated subjects were mechanically ventilated on a FlexiVent system (SCIREQ) at a rate of 150 breaths per minute with tidal volumes of 10 ml/kg and positive end-expiratory pressure of 3 cmH_2_O. Mice were then paralyzed using pancuronium bromide (1 mg/kg) via i.p. injection. Deep inflation and single-frequency forced oscillation maneuvers were run in triplicate on each subject to obtain parameters of inspiratory capacity (A), respiratory system resistance (Rrs), compliance (Crs) and elastance (Ers).

### House Dust Mite model of allergic airway inflammation

Mice were first sensitized with intranasal delivery of 1 μg of HDM extract (Greer Laboratories cat# XPB70D3A2.5) or PBS and were later challenged with 10 μg HDM extract for 4 consecutive days on days 7-10 post-sensitization. Analyses of ILC2s in lungs were performed 24 h after the final challenge.

### Single cell RNA sequencing (scRNA-seq) preprocessing and analysis

Female and male mice were infected with PR8 virus intranasally, and lungs were harvested at day 7 p.i. Lung ILCs were pooled from 4-5 female or male mice and sorted as lineage-negative (CD3, CD5, TCRβ, B220, CD11b, CD11c, TER119, Gr-1, CD19) CD90^+^IL-7Rα^+^ cells using a BD FACS Aria II instrument. Female and male ILCs were tagged with distinct Totalseq anti-mouse Hashtag (1 and 2) mAbs (anti-CD45 and MHCI) (Biolegend) to facilitate separation of the ILCs by sex in the sequencing analysis.

#### Single-cell multiplexing and sequencing

Cell capture was performed using the 10x Genomics Chromium system followed by sequencing on an Illumina HiSeq 3000, each according to the manufacturer’s protocol. Following qPCR quantification of the final library, each library was loaded onto a single lane of a HiSeq 3000 using read lengths of 28 bp for the first read, 98 bp for the second read, and an 8 base index read. Upon completion of sequencing the raw bcl files were processed using the 10x Genomics Cell Ranger (v3.0.2) informatics pipeline, using the mm10 murine reference genome. The protocol was repeated to produce two total cell pools, yielding transcriptome profiles for 4976 and 5516 cells for pool 1 and pool 2, respectively. scRNA-seq data were deposited in GEO as GSE190941.

#### Single-cell RNA-seq quality control and cell sex determination

We performed quality control on each pool separately by removing cells under any of the following conditions: 1) The number of expressed genes fell outside the 1^st^ or 99^th^ quantile across all cells in the pool; 2) The proportion of mitochondrial reads was above the 99^th^ quantile across all cells; 3) The sex of the donor mouse was not determined, or 4) the probability of being a cell doublet was estimated to be greater than 33% as estimated by the Scrublet method (61). This probability threshold was chosen based on the 99^th^ percentile of Scrublet estimates across all cells. 7388 total cells remained after quality control. Cell hashing antibody tags labeling male and female mouse cell pools were recovered from reads via native cellranger count functionality as well as the CITE-seq-count tool (https://cite-seq.com/). We also used the following scheme to determine cell donor sex: If the sum expression of Y-chromosome-specific genes Uty, Ddx3y and Eif2s3y was positive and zero Xist counts were observed, then the cell is male. Otherwise, if Xist expression is positive and the sum of the above Y-chromosome genes is zero, then the cell is female.

#### Assembly of pooled datasets and clustering

To control for technical variation across pools, we performed data integration using R package *Seurat* v3.0 (62). Pools were individually normalized using the *SCTransform* function with default options. Next, 5000 anchor features were selected across both pools and datasets were merged with the *IntegrateData* function. Linear dimensionality reduction was performed using PCA and retaining the first 90 principal components. Clustering was performed using the Louvain method via the *FindNeighbors* and *FindClusters* functions with the following options: dims=1:50, resolution=0.3.

#### Cell type determination and differential expression analysis

The projected single-cell data was embedded using uniform manifold approximate projection (UMAP, default parameters) for visualization. Gene markers unique to each cluster were obtained using the Seurat function *FindAllMarkers* and used to guide cluster identity. Sex-specific counts of each cell type are given in Fig. S4C. To determine differentially expressed genes by sex for a given cell cluster, we used the FindMarkers function in Seurat (options min.pct =0.25, logfc.threshold=0.25). We adjusted the statistical significance of each test for multiple hypotheses by controlling the false discovery rate (FDR) at 5%.

#### Cell type proportion analysis

To determine whether cell type (i.e. cluster) proportions changed between male and female mice, we modeled the number of cells as a random variable using Poisson regression. The R function *glm* was used to model cell counts as a function of sex and donor pool, including the total cells per pool as an offset. The *p-*value for the effect of the modeled regression coefficient for sex was obtained using a Wald test. To account for multiple hypotheses, these values were adjusted by controlling the FDR at 5%. Note that these tests are not independent and are not reflective of overall compositional differences.

#### Comparison to a published ILC2 dataset

These data were generated following the older version of the mapping and annotating query datasets vignette based on web-published protocols, https://satijalab.org/seurat/archive/v3.0/integration. Briefly the data sets from the Locksley (32) and Kovats groups were integrated, and anchor markers were identified between the two sets. The anchors were then transferred to the data from the Locksley group, which was used as the reference dataset. Using data from the Kovats group as a query, the Locksley data was separated into the 8 clusters identified by the Kovats group.

### Statistics

Comparison between two groups was performed using a two tailed unpaired t test or a Mann– Whitney U test using Prism 9 software. A one-way or two-way ANOVA followed by a Tukey or Sidak’s multiple comparison test was performed for multiple group comparisons. Weight loss curves were analyzed by a repeated measure 2 way ANOVA with mixed-effects analysis, followed by a Sidak’s multiple comparison test. A p value < 0.05 was considered significant, with *, p<0.05; **, p<0.01; ***, p<0.001; ****, p<0.0001. All graphs show mean and SD.

---

**Table S1.**
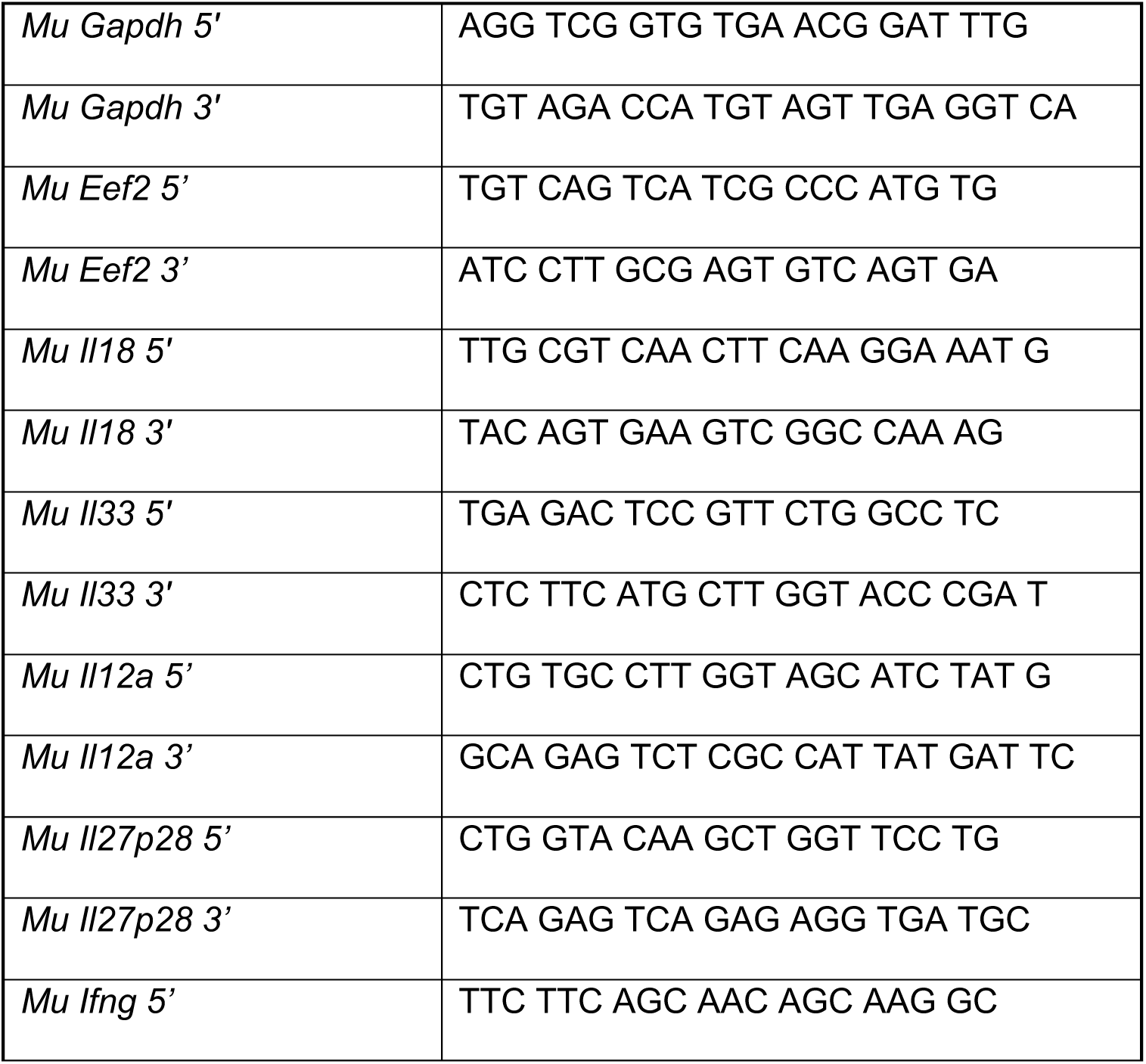
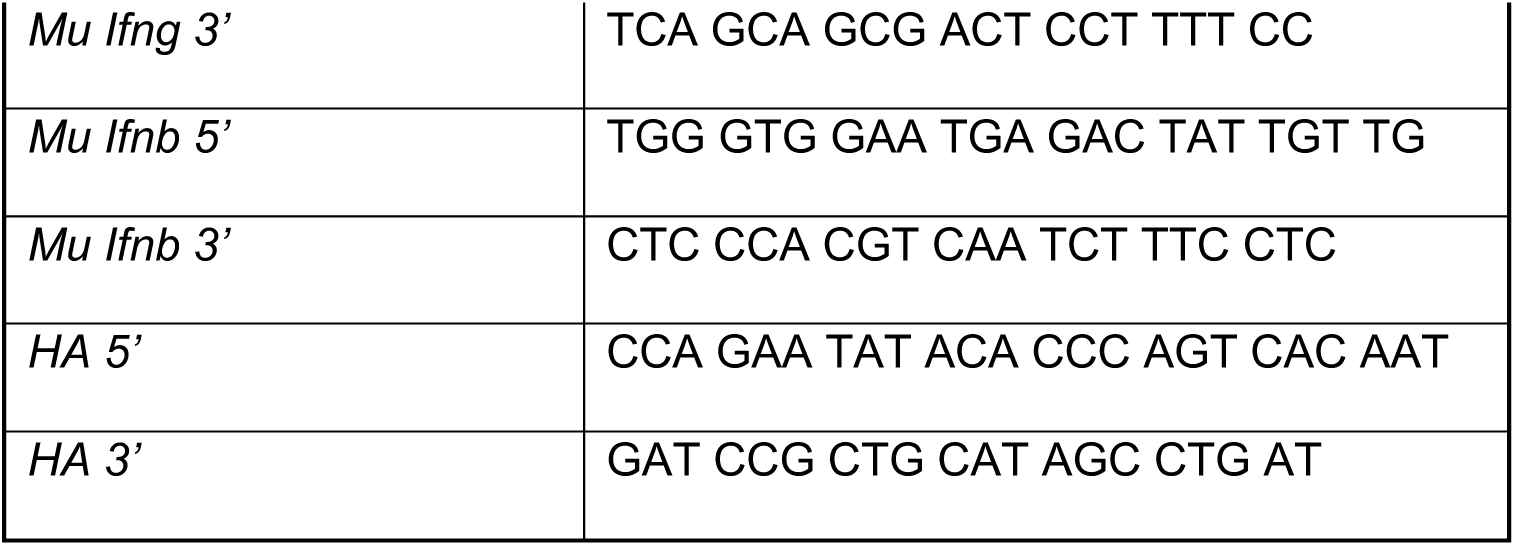
Primer Sequences for qPCR.

## Acknowledgements

We thank the staff of the OMRF Flow Cytometry Core, the Clinical Genomics Center, the Quantitative Analysis Core and the Comparative Medicine Department for expert technical help. This work was supported by NIH HL119501 and the Presbyterian Health Foundation (to S. Kovats), NIH AI129458 (to JA and HB) and by an OMRF Patricia and Don Capra Predoctoral Fellowship (to S. Kadel).

## Author contributions

S. Kadel designed the study, performed experiments, analyzed data and wrote the manuscript. AK, ST, RM, EA, IH, JG performed experiments and analyzed data. HB, JA and RP helped design the study and analyzed data. S. Kovats procured funding, designed and supervised the study, analyzed data, and wrote the manuscript.

---

**Fig. S1.**
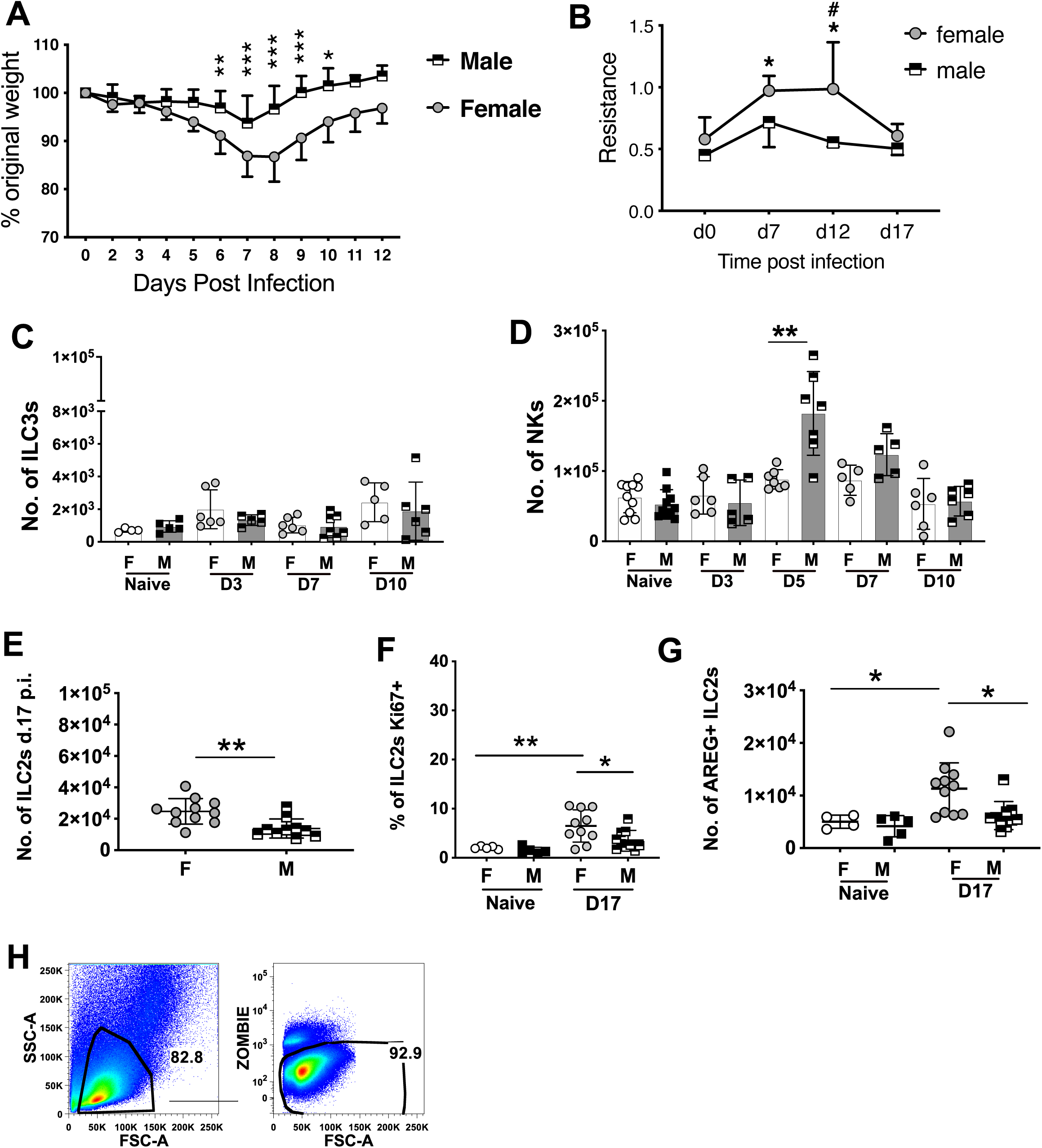
Supporting data for Fig. 1. (**A**) Weight loss curves of infected female and male mice over 12 days post-infection. Data were analyzed by a repeated measure 2 way ANOVA with mixed-effects analysis, followed by a Sidak’s multiple comparison test. Shown are mean +/- SD, n=12 of each sex. (**B**) Measurement of lung function as airway resistance on a Flexivent instrument on the indicated days post-infection. Data were analyzed using a 2 way ANOVA, n=3-4 per time point. Differences (indicated by *) within each sex over time relative to day 0 were assessed using a Tukey’s multiple comparison test. Differences (indicated by #) between each sex at each time point were assessed using a Sidak’s multiple comparison test. (**C**) Number of ILC3s (defined as defined as Lin^−^TCRβ^−^IL-7Rα^+^ RORGT^+^EOMES^−^) and (**D**) NK cells (defined as Lin^−^TCRβ^−^T-BET^+^EOMES^+^) in the lung at the indicated days post-infection. (**E**) Number of lung ILC2s day 17 p.i. (**F**) Fraction of ILC2s Ki67^+^ in mice naïve and day 17 p.i. (**G**) Number of AREG^+^ ILC2s in mice naïve and day 17 p.i. (**H**) Gating of viable lung cells (pre-gate to Fig. 1A). For panels C-G, significance was evaluated using Mann-Whitney or Student’s t tests at each time point (panels C,D,E) or a one way ANOVA with Tukey’s multiple comparison test (panels F,G).

**Fig. S2.**
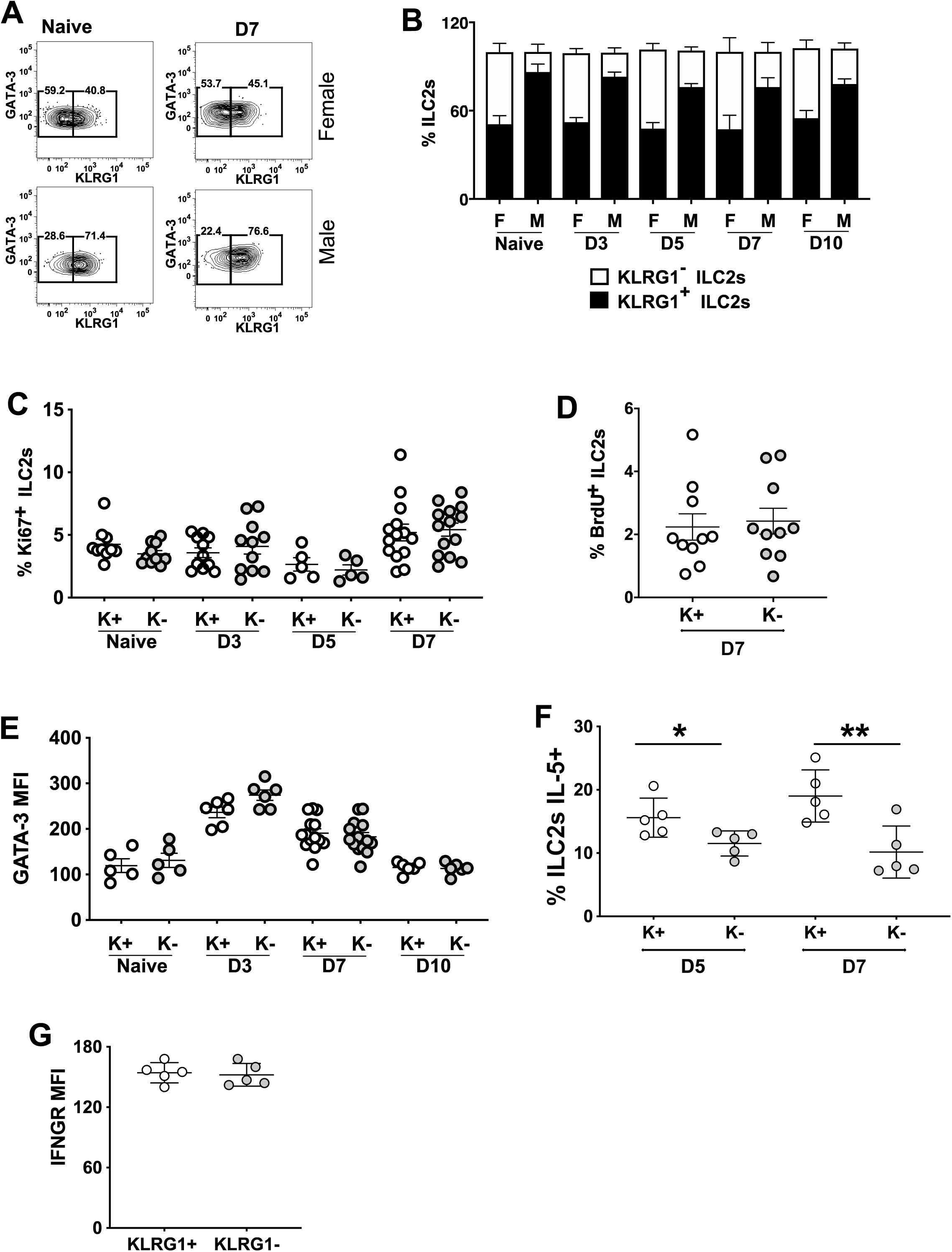
IAV infection does not alter the ratio or phenotype of KLRG1+ and KLRG1^−^ ILC2s in females or males. (**A**) Staining of KLRG1 on ILC2s in mice naïve or on day 7 p.i. (**B**) Relative proportion of KLRG1^+^ and KLRG1^−^ ILC2s in males and females on the indicated days p.i. Data represent 2-4 experiments per time point, with 4-9 mice per group. (**C**) Fraction of KLRG1^+^ (K^+^) and KLRG1^−^ (K^−^) female ILC2s that were Ki67^+^ at the indicated time points. (**D**) GATA3 MFI of KLRG1^+^ and KLRG1^−^ female ILC2s at the indicated time points. (**E**) Fraction of KLRG1^+^ and KLRG1^−^ female ILC2s that were BrdU^+^ at day 7 p.i. (**F**) Fraction of KLRG1^+^ and KLRG1^−^ female ILC2s that were IL-5^+^ at the indicated time points. (**G**) IFNGR MFI on KLRG1^+^ and KLRG1^−^ ILC2s in naïve female mice.

**Fig. S3.**
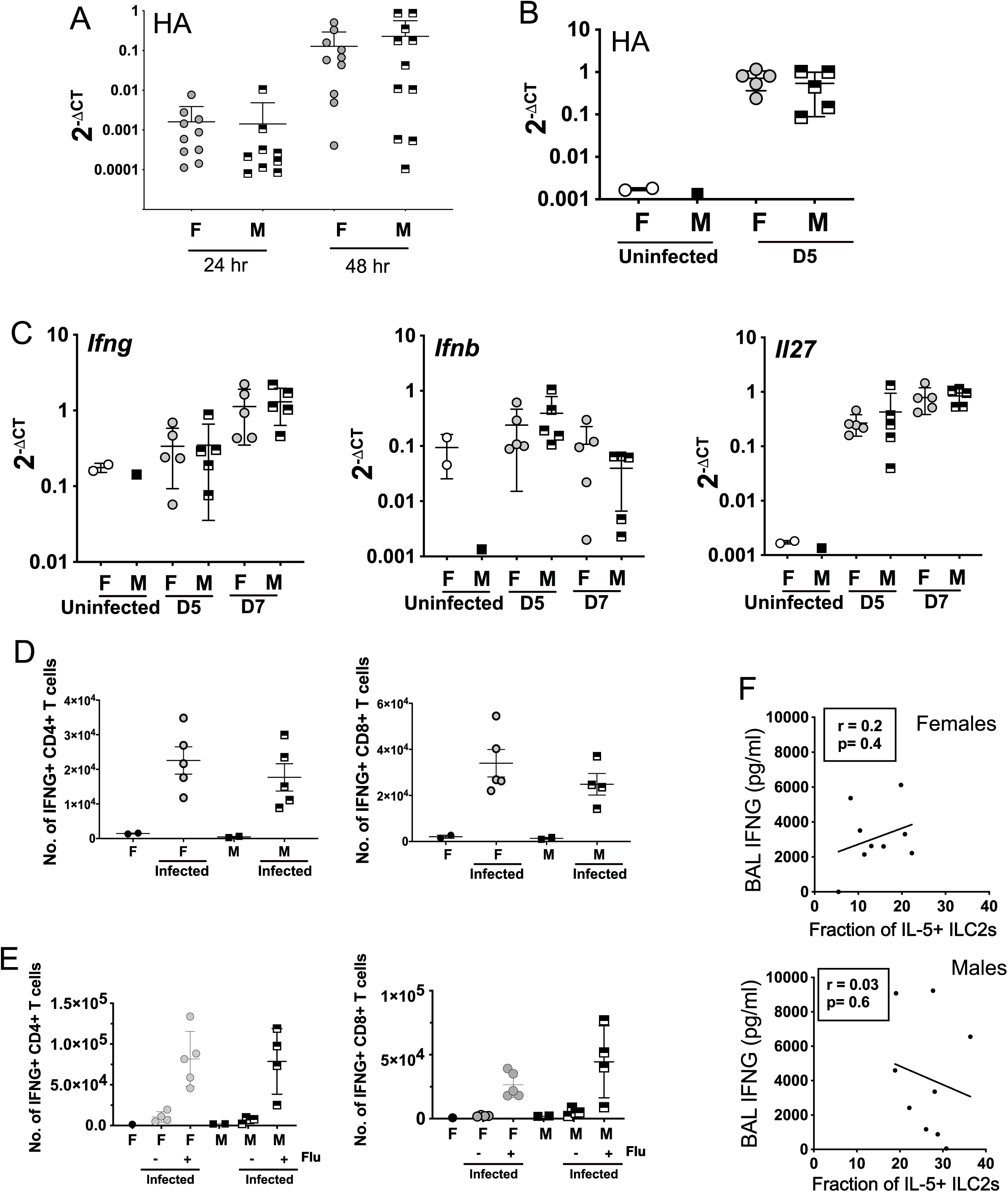
Supporting data for Fig. 3. (**A**) Quantification of viral HA RNA levels by qPCR in female and male lungs at 24 and 48 hr p.i. (**B**) Quantification of viral HA RNA levels by qPCR in female and male lungs at day 5 p.i. (**C**) Quantification of *Ifng, Ifnb and Il27* RNA levels by qPCR in female and male lungs at days 0, 5 and 7 hr p.i. (**D**) Numbers of total IFNG^+^ CD4^+^ and CD8^+^ T cells in lung on day 7 p.i. (**E**) Numbers of IFNG^+^ CD4^+^ and CD8^+^ T cells specific for influenza peptides in lung on day 12 p.i. (**F**) Correlation between IFNG levels in BAL with the number of IL-5^+^ ILC2s in lung in females and males.

**Fig. S4.**
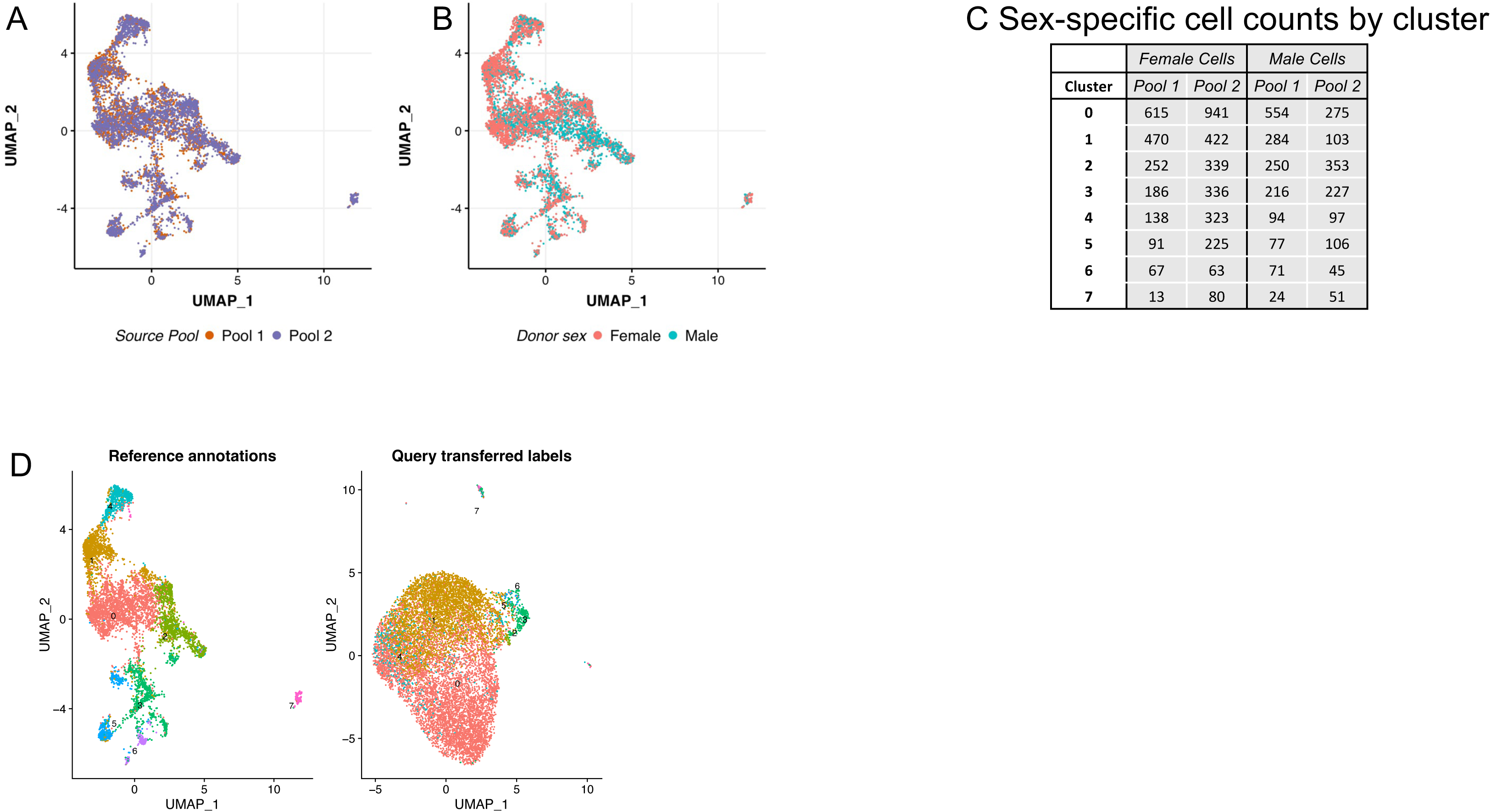
Supporting data for Fig. 5. (**A**) UMAP embedding of PCA projection, colored by dataset (pool 1 or 2). (**B**) UMAP embedding of PCA projection, colored by sex. (**C**) Sex specific cell counts by cluster in pools 1 and 2 of female and male cells. (**D**) Integration of our scRNA-seq clusters (reference annotation) with a dataset of combined male and female lung ILC2s in naïve mice (query transferred labels) [reference 32]. The 8 clusters in each dataset are color-coded.

**Fig. S5.**
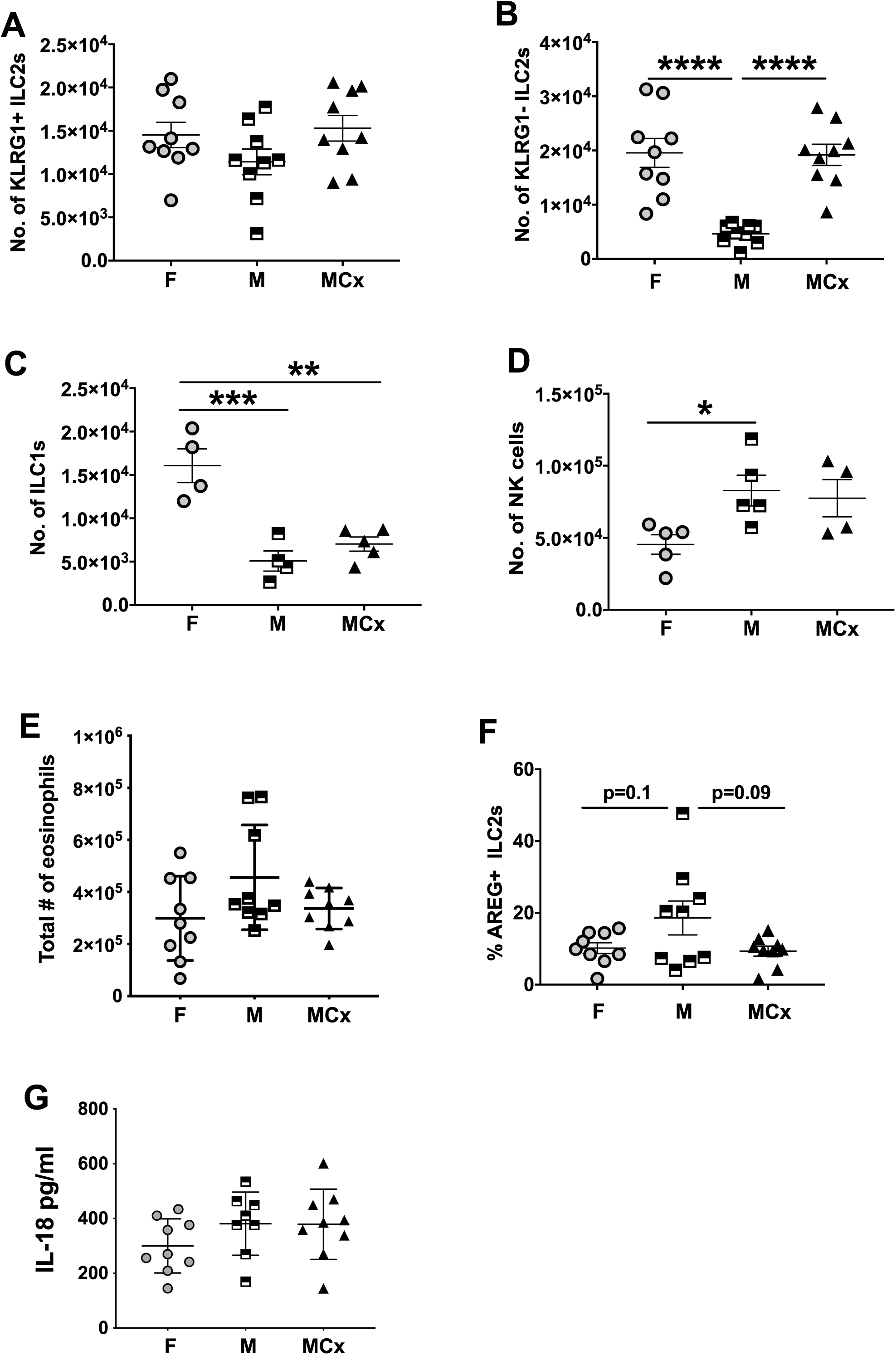
Supporting data for Fig. 6. Analyses of cell subsets in lungs of female, male and orchiectomized (MCx) males on d.7 p.i. with IAV. Numbers of (**A**) KLRG1^+^ and (**B**) KLRG1^−^ ILC2s. Numbers of ILC1s (**C**) and NK cells (**D**). (**E**) Number of eosinophils. (**F**) Fraction of ILC2s AREG^+^. (**G**) Levels of IL-18 in BAL.

**Fig. S6.**
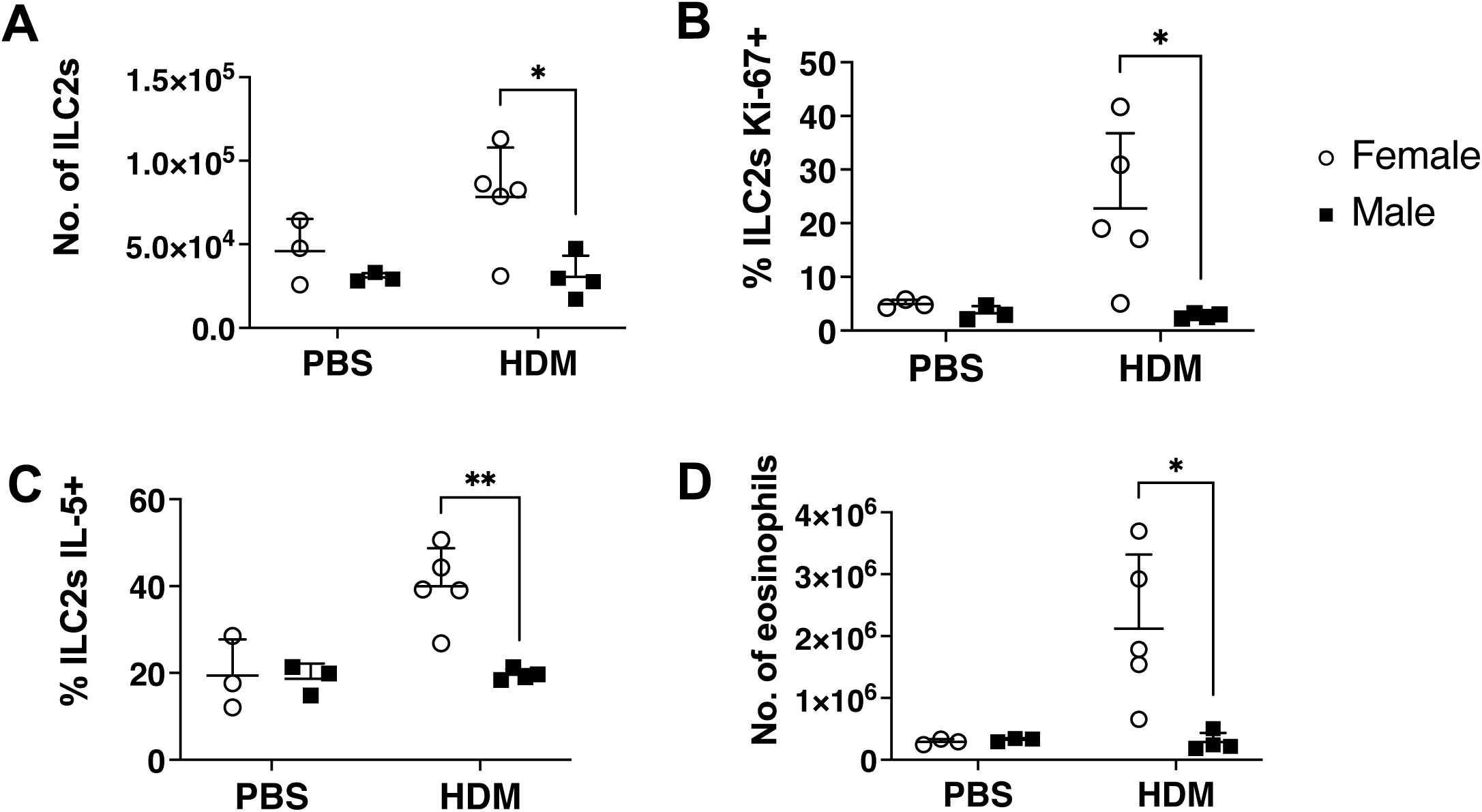
Sex differences in ILC2s in a model of House Dust Mite (HDM) challenge inducing allergic airway inflammation. Mice were sensitized with intranasal delivery of 1 μg of HDM extract or PBS and were later challenged with 10 μg HDM extract for 4 consecutive days on days 7-10 post-sensitization. Analyses of ILC2s in lungs were performed 24 h after the final challenge. (**A**) Number of ILC2s in males or females. (**B**) Fraction of ILC2s Ki67^+^. (**C**) Fraction of ILC2s IL-5^+^. (**D**) Number of eosinophils. Data were analyzed using a two-way ANOVA comparing sex and treatment with a Sidak’s multiple comparison test.

